# Engineering tissue-mimetic hydrogels with continuous viscoelastic gradient for programmed cell migration

**DOI:** 10.1101/2024.12.24.630289

**Authors:** Haochen Yang, Ziyuan Li, Meng Li, Yuxing Shang, Linjie Chen, Yifan Ge, Junji Zhang, He Tian

## Abstract

The dynamic, viscoelastic nature of the extracellular matrix (ECM), with variations in stiffness and viscosity, plays a key role in cellular behaviors like mechano-sensing, migration, and force generation. Viscoelastic hydrogels are important for mimicking the ECM to study cell migration and tissue engineering. However, creating hydrogels with continuous viscoelastic gradients is challenging due to issues with spatial resolution, reproducibility, and accurately replicating native tissue properties. Here we present a novel approach to generate hydrogels with precisely controllable viscoelastic gradients using drop-by-drop condensation. This approach allows fine-tuned replication of the tissue-specific local mechanical properties, resulting in hydrogels of heterogeneous viscoelasticity. Directed migration and separation of multiple cell lines are thus driven by local viscosity and elasticity rather than solely stiffness. By incorporating non-linear mechanical gradients, this technique provides insights into ECM viscoelasticity’s role in cellular behavior and offers a versatile platform for advanced tissue engineering and regenerative medicine applications.

## Introduction

The mechanical properties of biological tissues exhibit viscoelastic behaviors arising from complex interactions between cells and their surrounding extracellular matrix (ECM)^1,2^. Within individual cells, filamentous actin and myosin II form the actomyosin cortex and contractile filaments, enabling tension generation and bidirectional force transduction between the cell and the ECM^3,4^. This dynamic machinery allows cells to sense and interpret mechanical signals from the ECM, reorganize their actomyosin structure, remodel the ECM, initiate migration, and recalibrate intercellular communication, ultimately influencing their fate^4,5^. The mechano-interaction between the ECM and cells, along with cellular migration behavior, plays a critical role in numerous physiological and pathological processes, such as embryogenesis, wound healing, cancer invasion, tissue repair, and homeostasis^1,2,4,6^.

Native ECM is a meshwork of fibrous proteins and polysaccharides, with spatial-temporal variation of their chemical properties, compositions, and interactions^10,11^, resulting in a non-linear property that cannot be easily replicated using typical hydrogels^7,12-14^. Focal adhesions (FA), the major mechano-linker between ECM and actomyosin networks forms the classical molecular clutch that is responsive to the stiffness of ECM and determines cell migration^5,7^. In this model, cells cultured on soft ECMs tend to exhibit weaker adhesions and reduced contractility, while stiffer ECMs promote the development of more stable and robust adhesions, enhancing contractile responses and causing the durotaxis behavior to promote cell migration toward stiffer substrate^3,5^. Notably, recent studies indicate that the viscosity of the local environment also impacts cellular mechano-sensing^8^. Yet, it remains unclear how viscosity and elasticity contribute together in cell mechano-sensation and the following behaviors, especially under a biomimetic nonlinear alternation of the mechano-properties.

Engineered hydrogels with dynamic structures have been deeply investigated and applied to programmed cell and organoid culturing as interactive “living” material^15,16^. Significantly, new insights into these dynamic viscoelastic hydrogels would provide general strategies for the rational design of biomimetic ECMs^17^, which are indispensable and applicable in cell mechanical regulation and tissue engineering. To this end, fabricating an ECM-mimicking hydrogel system that allows gradient change of viscoelastic properties is an ideal tool for investigating force generation and related cell biology mechanisms^18^. Current methods for fabricating gradient hydrogels typically involve the copolymerization of two gel formula droplets placed on either side of a coverslip^19^. While this approach is straightforward, it often leads to hydrogels with unpredictable mechanical gradients and inadequate spatial resolution, hindering the precise customization and replication of real tissue mechanics within a single gel. Although photo-polymerization offers the advantage of enhanced spatiotemporal resolution in gradient hydrogel preparation^20^, it still falls short of mimicking the intricate multi-gradient structures observed in native tissues. Moreover, these existing techniques primarily focus on creating elastic gradients in stiffness, neglecting the more complex viscoelastic gradients crucial for accurately matching tissue properties.

In this study, we present a novel strategy for engineering hydrogels that achieve stable and reproducible viscoelastic gradients, closely mimicking the mechanical properties of various tissues that allows quantitative characterization of local mechanical properties. Based on this platform, we created cell-culture substrates featuring heterogeneous mechanical characteristics, enabling the programmed migration and separation of multiple cell lines through mechano-transduction on a single hydrogel. Our experiments demonstrated that distinct types of cells, including chondrocytes, NIH-3T3 and C2C12, could exhibit directed migration in response to substrate viscoelasticity (Fig. 1a). Notably, our findings challenge traditional notions of traction force behavior on elastic substrates. We observed that cellular contractile responses are not linearly correlated with substrate stiffness. When viscosity is considered, maximum traction forces occurred on matrices that closely matched the mechanical properties of the originating tissues (Fig. 1b and 1c). Further analysis at the single-cell level revealed that focal adhesion maturation is also influenced by changes in viscosity independently from intercellular connections. In this way, we showed that our innovative approach can program guided migration and effectively sort different cell clusters within one system, which allows more delegated chemical-based tissue engineering in the future.

**Fig. 1:**
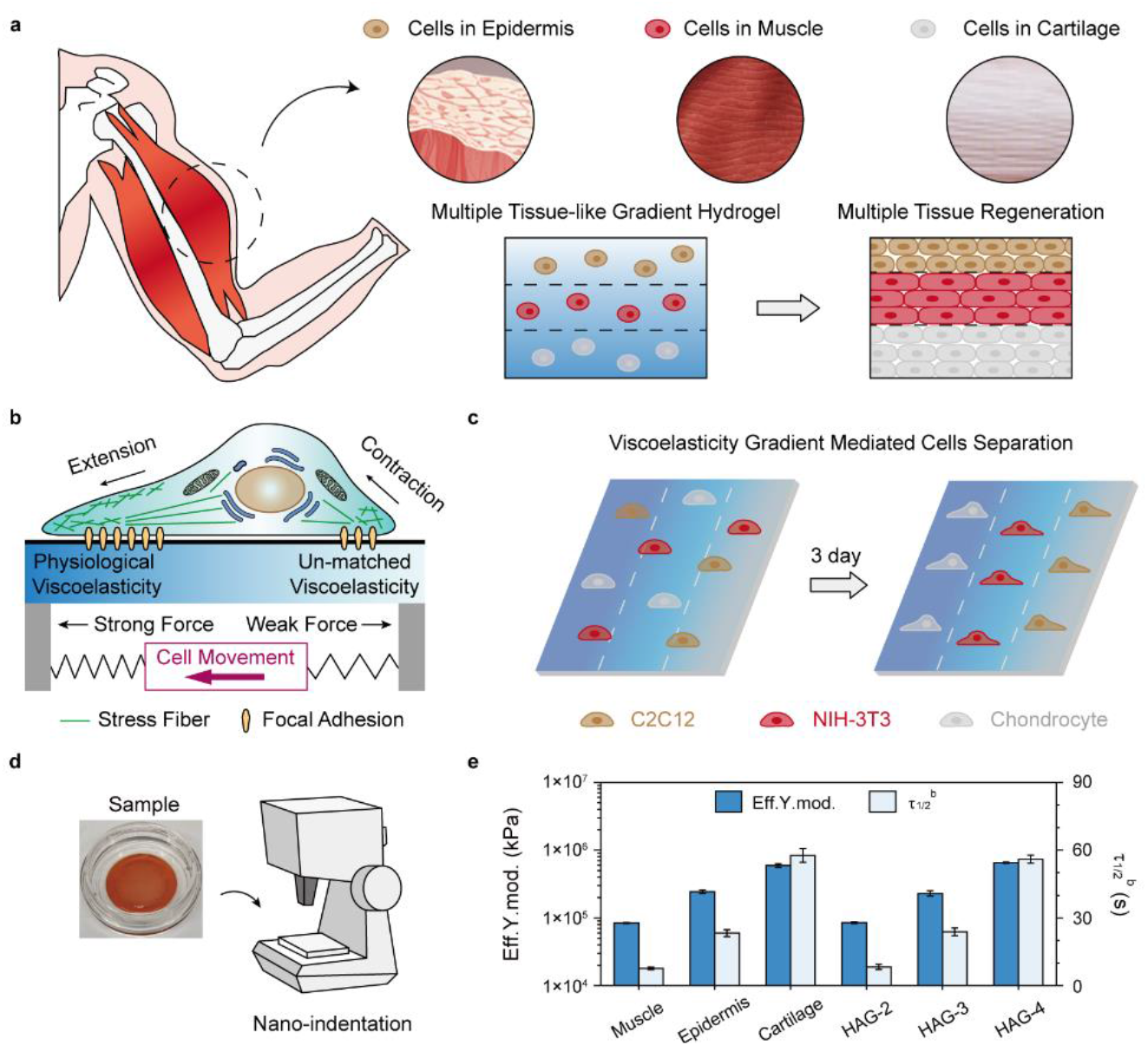
Directed migration of multiple cells by viscoelastic gradient hydrogels based on mechano-transduction. a) Multiple tissue-like hydrogel for multi-tissue regeneration. b) Cell located on viscoelastic gradient substrates form different numbers of focal adhesions at the front and back ends, leading to asymmetric traction and cell polarization, which in turn guides directed migration. c) Diagram of mixed cells separation mediated by gradient viscoelastic hydrogel. d) Nano-indentation device for gradient viscoelasticity measurement. e) Effective Young’s modulus and τ_1/2_^b^ (measured by nano-indentation) of muscle, epidermis, cartilage, **HAG-2, HAG-3** and **HAG-4**.

## Results

### Engineering tissue-mimetic hydrogel with precisely controlled gradient viscoelasticity

To replicate the viscoelastic properties of native extracellular matrices (ECMs) with precisely defined viscoelasticity, we developed synthetic hydrogels (**HAG**) composed of oxidized hyaluronic acid and gelatin copolymer networks (Fig. 2a and Supplementary Fig. 1). The flexible natural polysaccharides impart weak dynamic interactions through hydrogen bonding and the release of entanglements. The dynamic covalent imine bonds generated within the network, on the other hand, provide energy dissipation (stress relaxation) and enhanced mechanical strength (stiffness). Inspired by natural ECMs, this strategy allows for tunable viscoelasticity by adjusting the ratios of the two flexible chains, thereby balancing weak and dynamic solid interactions. Furthermore, gelatin introduces binding sites (RGD) that facilitate interactions between the hydrogel network and cells^21,22^. Fourier Transform Infrared Spectroscopy (FT-IR) analysis confirmed the successful formation of **HAG** hydrogels, where characteristic peaks at 1646 and 1544 cm^−1^, corresponding to C=N bonds formed through the condensation reaction between the aldehyde groups (-CHO) of oxidized hyaluronic acid and the amino groups (-NH_2_) of gelatin (Supplementary Fig. 2). To acquire the most effective hydrogel formula to mimic targeted tissues, the viscoelastic parameters, mechanical strength (stiffness; G’) and stress relaxation time (τ_1/2_), of acellular cartilage, epidermis, and muscle were first determined through rheology measurements (Table 1 and Supplementary Fig. 3) and adopted as the interpolation to establish a hydrogel formulation library (Fig. 2b-2c, Supplementary Fig. 4-8 and Supplementary Table 1; See Methods for details). Three hydrogel formulae with different oxi-HA/gelatin ratio (**HAG-2** to **HAG-4**, Table 1) were tested to match with corresponding acellular muscle, epidermis and cartilage tissues, respectively. Two additional hydrogel formulae, **HAG-1** and **HAG-5** with stiffer and softer mechanics, respectively, were also obtained to fabricate gradient viscoelastic hydrogels with above tissue-related **HAG** formulae (Supplementary Fig. 9 and Table 1; see Methods for details).

**Table 1.** Injection concentration of different hydrogels and viscoelastic parameter of muscle, epidermis, cartilage and different hydrogels. G’ and τ_1/2_^a^ were determined by rheometer. Effective Young’s modulus and τ_1/2_^b^ were determined by nano-indentation.

| | oxi-HA<br>($\omega/v$) | gelatin<br>($\omega/v$) | G'<br>(Pa) | $\tau_{1/2}^a$<br>(s) | Eff. Y.<br>mod.<br>(kPa) | $\tau_{1/2}^b$<br>(s) |
| --- | --- | --- | --- | --- | --- | --- |
| muscle |  |  | 657 | 225 | 84 | 8 |
| epidermis |  |  | 970 | 302 | 244 | 23 |
| cartilage |  |  | 2014 | 413 | 594 | 58 |
| HAG-1 | 1.8 % | 21.2 % | 420 | 143 | 18 | 3 |
| HAG-2 | 3.8 % |  | 649 | 226 | 85 | 8 |
| HAG-3 | 7.8 % |  | 973 | 303 | 230 | 24 |
| HAG-4 | 14.8 % |  | 2077 | 411 | 654 | 56 |
| HAG-5 | 16.0 % |  | 2538 | 436 | 856 | 98 |

**Fig. 2:**
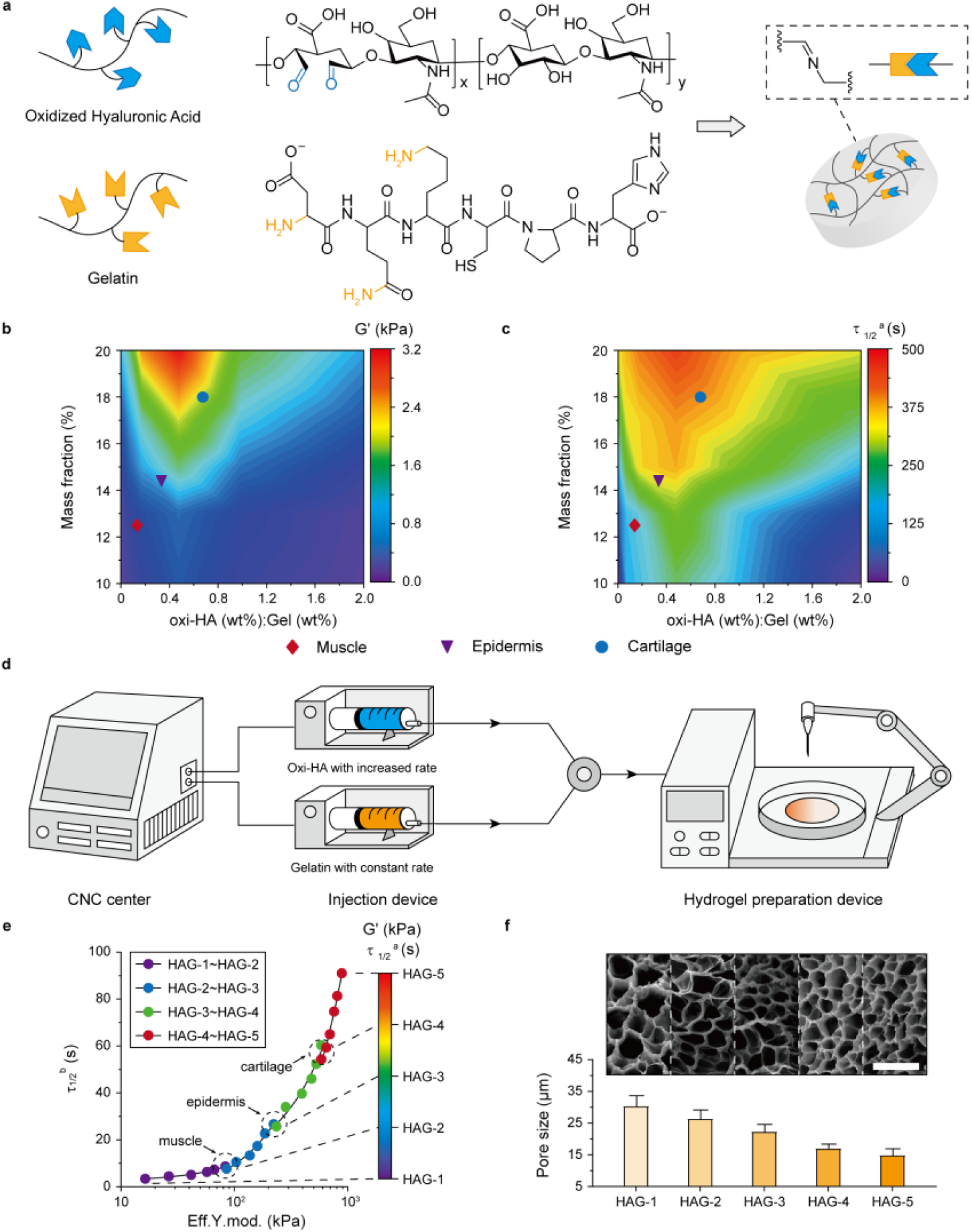
Construction of the multiple tissue-like gradient hydrogel. a) Construction of viscoelastic hydrogel by oxi-HA and gelatin. b) G’ and c) τ_1/2_^a^ (measured by rheometer) heat map of hydrogel formulation library. d) Technological process for preparing gradient viscoelastic hydrogel. e) The effective Young’s modulus and τ_1/2_^b^ of gradient hydrogels characterized by nano-indentation. f) SEM images and pore size analysis of hydrogels. Scale bar = 30 μm.

Biomimetic **HAG** hydrogels with viscoelasticity gradient were developed by meticulously preparing pre-gelation droplets with varied proportions of two flexible chains. Hydrogel precursor droplets containing gelatin (0.212 g/mL) and oxi-HA (0.018 g/mL – 0.160 g/mL) with tailored concentration gradients. The gradient hydrogel was then fabricated drop by drop through a syringe pump (SP-2000, Ningbo Anno Medical Equipment Technology Co., Ltd; *v*_*HA*_ = *v*_*gel*_ = 0.6 mL/min), enabling the formation of a continuous viscoelastic gradient between each droplet during gelation or copolymerization (Fig. 2d), while a robotic arm was applied to ensure uniform distribution of the gel by controlling the flow output. A computational numerical control (CNC) platform was employed to adjust the output formulations for subsequent droplet preparation to enhance reproducibility in the gelation process. This approach allowed for the creation of an orderly and customizable viscoelastic gradient within a single hydrogel. It should be noted that, due to small volume of each droplet (0.6 mL) and the precise control of viscoelasticity by adjusting the input ratio of oxi-HA to gelatin, we can simulate a continuous viscoelastic gradient similar to that of natural physiological tissues (Fig. 2e).

The gradient viscoelasticity (effective Young’s modulus, E; stress relaxation time τ_1/2_) in **HAG** hydrogel was assessed using nano-indentation techniques (Fig. 2e, Table 1 and Supplementary Fig. 10). To identify the tissue-related viscoelastic regions and standardize the mechanical data for the following experiments, the effective Young’s modulus and stress relaxation of targeted tissues were also characterized by nano-indentation (Fig. 1d-1e and Table 1). For each target tissue, three regions are randomly selected, and multiple tests are performed within each region. The average viscoelasticity (effective Young’s modulus, E; stress relaxation time τ_1/2_) is calculated and served as the viscoelasticity replication index. As illustrated in Figure 2e, the hydrogel displayed a continuous viscoelasticity gradient in both effective Young’s modulus (18 kPa - 856 kPa) and relaxation (3 s - 98 s) profiles, with corresponding viscoelastic parameters of measured tissues installed in respective formulae locations (**HAG-2** to **HAG-4**; Fig. 2e and Supplementary Fig. 10). Corresponding scanning electron microscopy (SEM) images showed a linear decrease in pore size from 30.2 μm to 14.7 μm, transitioning from the softer, faster-relaxation **HAG-1** to the stiffer, slower-relaxation **HAG-5** (Fig. 2f). This observation is in excellent agreement with the mechanical property measurements. Hence, a tissue-benchmarked hydrogel with precisely controlled viscoelasticity gradient was successfully fabricated and engineered, mimicking both the heterogeneous and continuous mechanical features of actual tissue ECMs. Importantly, maintaining a constant gelatin ratio throughout the hydrogel series ensures consistent cell adhesion biochemically via integrin-binding ligands (e.g., RGD), effectively mitigating any independent influence on cell migration (haptotaxis)^23-25^.

### Tissue viscoelasticity directs cell migration on viscoelastic gradient hydrogels

To further investigate cellular behavior on gradient hydrogels, we applied C2C12, NIH-3T3, and chondrocyte as the model cell systems. We prepared gradient gels with viscoelastic properties that correspond with the native tissue of the cell lines derived from. The cell survival and growth in viscoelastic gradient hydrogels were first evaluated on selected cell lines for at least 7 days (Supplementary Fig. 11), indicating good biocompatibility of fabricated hydrogels.

Each type of cell was seeded in the viscoelastic gradient regions of the hydrogels. After cell spreading, individual cell migration trajectories were recorded over a 12.5-hour period^26^. C2C12 cells displayed a directional movement toward the muscular viscoelastic region (**HAG-2**), characterized by an effective Young’s modulus of 85 kPa and a stress relaxation time (τ_1/2_) of 8.3 s. This region is identified as stiffer compared to the softer **HAG-1** region (17 kPa, τ_1/2_ = 2.9 s) and softer than the epidermis edge represented by **HAG-3** (230 kPa, τ_1/2_ = 23.8 s) (Fig. 3a (i) and (ii), Fig. 3b (i) and (ii)). In contrast, in control experiments where homogeneous hydrogels were applied with muscular viscoelasticity (**HAG-2**), C2C12 cells seeded on the hydrogel exhibited a directionless migration (Supplementary Fig. 12a-b), confirming a viscoelasticity-gradient induced directional cell migration on a physiological mimetic medium.

**Fig. 3:**
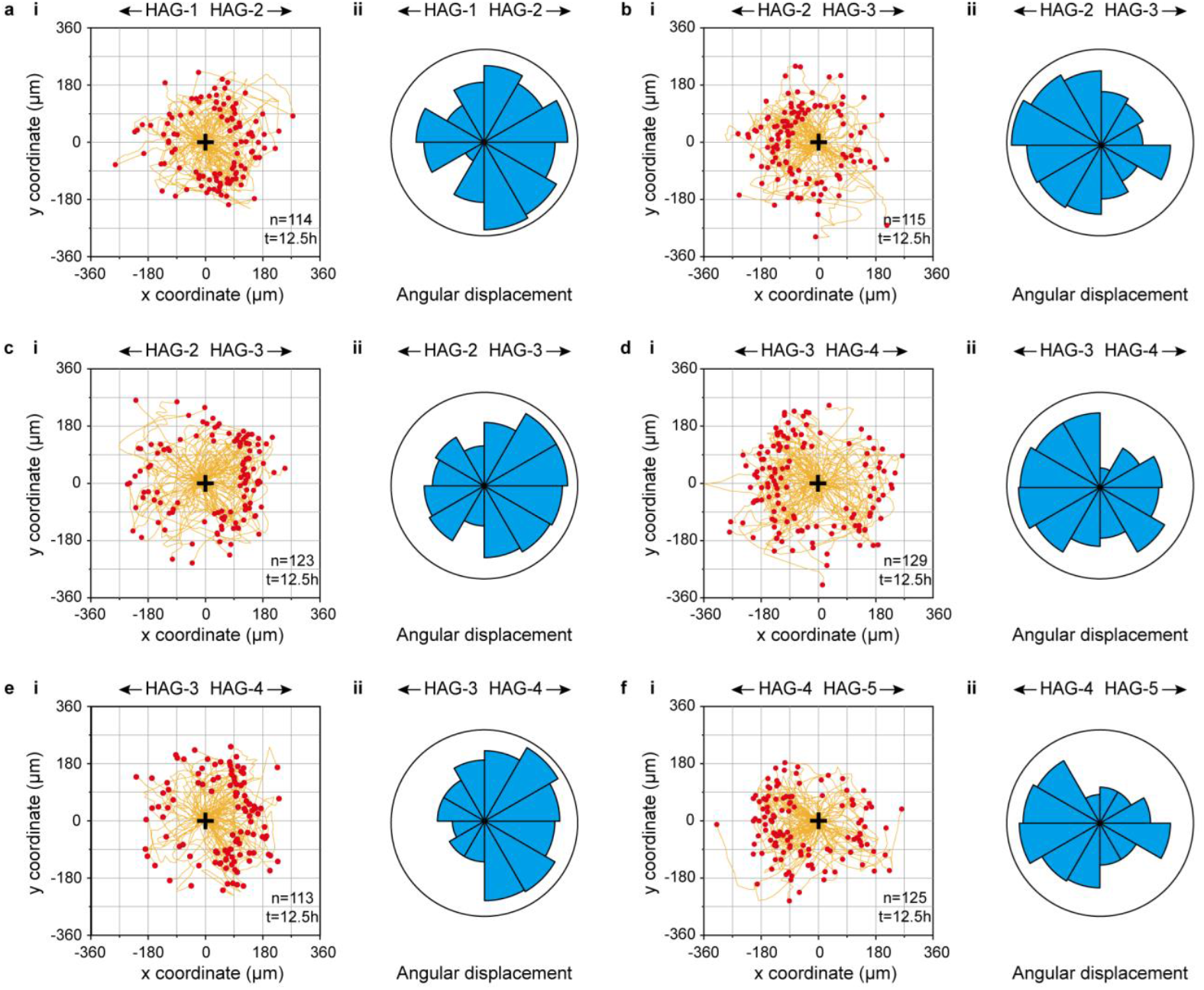
Cell migration trajectory. a) i) Migration trajectory of C2C12 on gradient **HAG-1∼HAG-2** viscoelastic regions, n = 114. ii) Angular displacement of C2C12 on gradient **HAG-1∼HAG-2** viscoelastic regions. b) i) Migration trajectory of C2C12 on gradient **HAG-2∼HAG-3** viscoelastic regions, n = 115. ii) Angular displacement of C2C12 on gradient **HAG-2∼HAG-3** viscoelastic regions. c) i) Migration trajectory of NIH-3T3 on gradient **HAG-2∼HAG-3** viscoelastic regions, n = 123. ii) Angular displacement of NIH-3T3 on gradient **HAG-2∼HAG-3** viscoelastic regions. d) i) Migration trajectory of NIH-3T3 on gradient **HAG-3∼HAG-4** viscoelastic regions, n = 129. ii) Angular displacement of NIH-3T3 on gradient **HAG-3∼HAG-4** viscoelastic regions. e) i) Migration trajectory of chondrocytes on gradient **HAG-3∼HAG-4** viscoelastic regions, n = 113. ii) Angular displacement of chondrocytes on gradient **HAG-3∼HAG-4** viscoelastic regions. f) i) Migration trajectory of chondrocytes on gradient **HAG-4∼HAG-5** viscoelastic regions, n = 125. ii) Angular displacement of chondrocytes on gradient **HAG-4∼HAG-5** viscoelastic regions.

Similarly, NIH-3T3 fibroblasts performed a directed migration towards their native epidermis viscoelastic regions (**HAG-3**) when seeded on gradient **HAG** hydrogel (Fig. 3c, d), rather than to the softer muscular side (**HAG-2**) or the stiffer cartilage side (**HAG-4** for cartilage viscoelasticity, Eff. Y. mod = 654 kPa and τ_1/2_ = 56 s), respectively. For chondrocytes, programmed cell migration towards native cartilage viscoelastic medium (**HAG-4**) was obtained when seeded on gradient **HAG** hydrogel, Fig. 3e, f, with limited migration to either softer epidermis side (**HAG-3**) or stiffer **HAG-5** edge (Eff. Y. mod = 856 kPa and τ_1/2_ = 98 s), respectively. Neither NIH-3T3 (Fig. 3c (iii) and Supplementary Fig. 12c, d) nor chondrocytes (Supplementary Fig. 12e, f) displayed evident directed migration in homogeneous hydrogels with epidermis (**HAG-3**) and cartilage (**HAG-4**) viscoelasticity, respectively. The above results suggested the directed migration behavior of C2C12, NIH-3T3, and chondrocytes could be regulated by programming the viscoelastic gradient of ECM mimetic medium, and the tissue viscoelasticity takes the preference in cell migration directions.

### Cells exert maximum traction forces on hydrogel gradients with tissue-matched viscoelasticity

Previous studies have shown that the local contractility of cells directs their migration and speed in response to ECM properties^27,28^. Both cellular traction and spreading are usually linear and correspond to the stiffness of the local environment^29^.Since cell migration in our gradient gels is not strictly durotaxis, we further investigated cell spreading and traction on **HAG**s. The spreading area of C2C12, NIH-3T3, and chondrocytes in different gradient **HAG** all displayed distinct morphological differences in response to the hydrogel viscoelasticity (Fig. 4a, i-iii). Each type of cell exhibited maximal spreading area when incubated on the hydrogels with corresponding physiological viscoelasticity. C2C12 exhibited a maximum average spreading area of 1166 µm^2^ on **HAG-2** comparing to on both stiffer (955 µm^2^; **HAG-3**) and softer (808 µm^2^; **HAG-1**) **HAG**s (Fig. 4b, i). Similar results were obtained in NIH-3T3 and chondrocytes. NIH-3T3 with a maximum average spreading area of 1164 µm^2^ was observed when plating on the **HAG-3** region with epidermis viscoelasticity (Fig. 4b, ii), compared to 845 µm^2^ and 937 µm^2^ plating on **HAG-2** and **HAG-4** regions, respectively. For chondrocytes, cells incubated on **HAG-4** with cartilage viscoelasticity showed a maximum spreading area of 941 µm^2^. On the contrary, cells seeded on either **HAG-3** and stiffer **HAG-5** regions presented less efficient spreading of 602 µm^2^ and 760 µm^2^, respectively (Fig. 4b, iii). The above results suggested that physiological tissue-like viscoelastic environments favor cells to perform the most efficient spreading when incubated on a native ECM-mimicking medium.

**Fig. 4:**
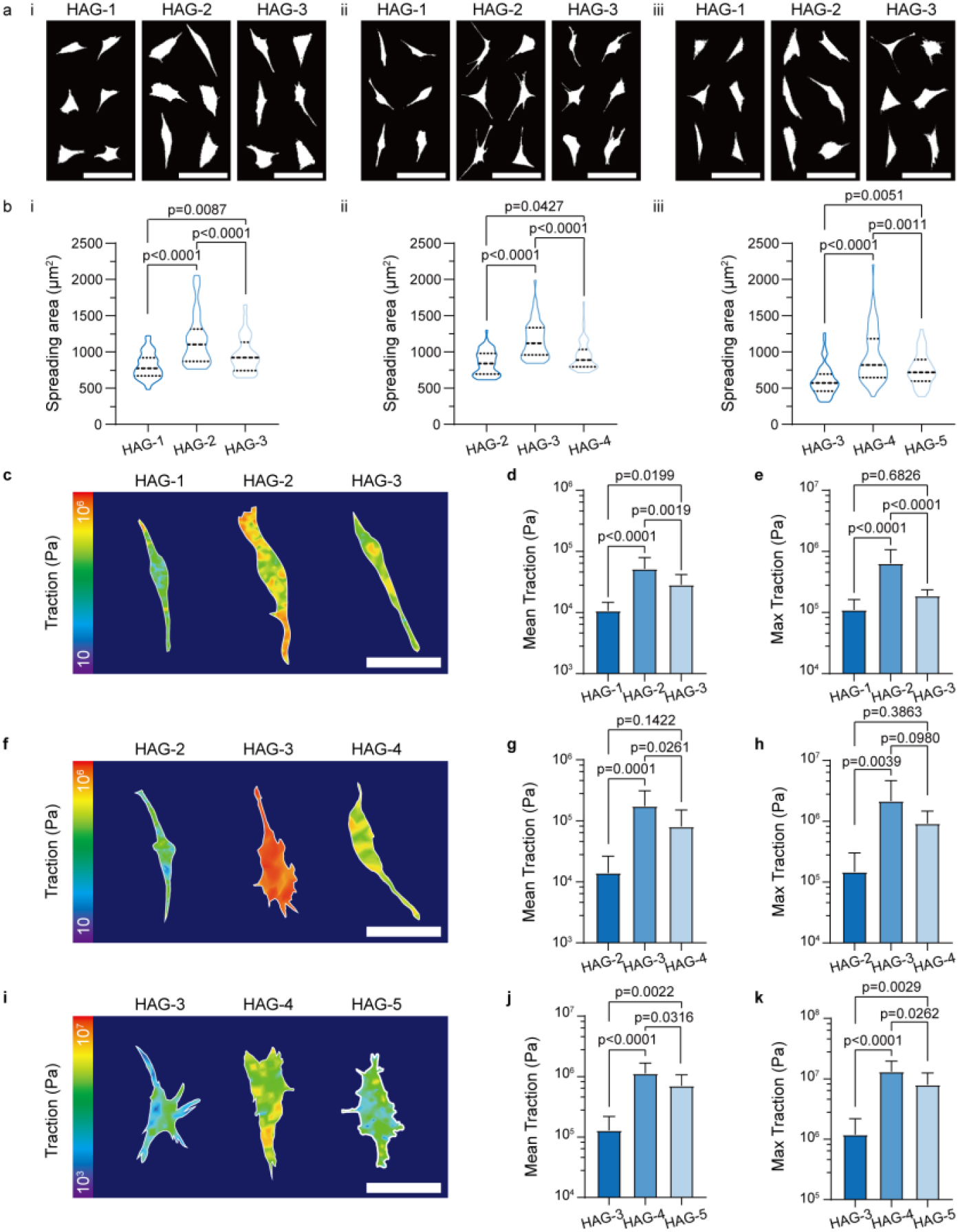
Cell traction force characterization. a) Morphology of i) C2C12 on **HAG-1, HAG-2** and **HAG-3**, ii) NIH-3T3 on **HAG-2, HAG-3** and **HAG-4**, and iii) chondrocytes on **HAG-3, HAG-4** and **HAG-5**, respectively. Scale bar = 50 μm. b) Quantitative analysis of spreading areas of i) C2C12, ii) NIH-3T3 and iii) chondrocytes, respectively. Ordinary one-way ANOVA was used for analysis of the data; n = 60 cells. Traction characterization of c) C2C12 on **HAG-1, HAG-2** and **HAG-3**, f) NIH-3T3 on **HAG-2, HAG-3** and **HAG-4**, and i) chondrocytes on **HAG-3, HAG-4** and **HAG-5**, respectively. Scale bar = 20 μm. Quantitative analysis of mean traction characterization of d) C2C12 on **HAG-1, HAG-2** and **HAG-3**, g) NIH-3T3 on **HAG-2, HAG-3** and **HAG-4**, and j) chondrocytes on **HAG-3, HAG-4** and **HAG-5**, respectively. Error bars, standard error of the mean (mean ± s.e.m). Ordinary one-way ANOVA was used for analysis of the data; n = 15, 13 and 12 cells, respectively. Quantitative analysis of maximal traction characterization of e) C2C12 on **HAG-1, HAG-2** and **HAG-3**, h) NIH-3T3 on **HAG-2, HAG-3** and **HAG-4** and k) chondrocytes on **HAG-3, HAG-4** and **HAG-5**, respectively. Error bars, mean ± s.e.m. Ordinary one-way ANOVA was used for analysis of the data; n = 15, 13 and 12 cells, respectively.

Traction force microscopy of C2C12, NIH-3T3, and chondrocytes seeded on corresponding viscoelastic gradient hydrogels was conducted (Figure 4c, 4f and 4i) according to previous reports^29^. Quantitative analysis showed C2C12 generated average traction of 51 kPa (me.t.f.) and 634 kPa maximum (ma.t.f.) on **HAG-2** (Fig. 4d, 4e), compared to cells incubated on hydrogel regions with “softer” **HAG-1** (10 kPa (me.t.f.)/110 kPa (ma.t.f.)) and on “stiffer” **HAG-3** (28 kPa (me.t.f.)/187 kPa (ma.t.f.)). NIH-3T3 and chondrocytes showed similar trends. NIH-3T3 displayed significantly larger mean traction force (175 kPa; Fig. 4g) and maximum traction force (2157 kPa; Fig. 4h) on epidermis benchmarked **HAG-3** viscoelastic region. In comparison, decreased values were revealed on the other two viscoelastic regions, respectively (Fig. 4g, 4h). Both mean (1125 kPa) and maximum (13168 kPa) traction forces also reached the peak values for chondrocytes seeded on cartilage mimicking **HAG-4** viscoelastic region (Fig. 4j, 4k), while falling sharply on viscoelastic regions deviating from the physiological mechanical microenvironment. These results revealed the tissue-mimicking viscoelastic microenvironment could promote native cells to generate larger traction forces. As the asymmetric traction force leads to cell polarization and propels cells to migrate towards the direction of higher traction force^30-33^, the above results might explain the physiological viscoelasticity-directed cell migrations on viscoelastic gradient hydrogels.

### Cell mechano-signaling correlates with the directed cell migration

In our migrating analysis, most cells were analyzed at single cell level, without cell-cell adhesion involved within our system. We hypothesized that the molecular clutch machinery could be one of the dominant mechano-sensation and transduction machinery^5,34-39^. In adhesive cells, receptors, primarily integrins, on the plasma membrane form connections between the extracellular matrix (ECM) and actomyosin networks through structures known as focal adhesions (FAs). Previous researches reveal that FAs, comprised of proteins such as integrins, focal adhesion kinases (FAKs), paxillin, talin, and vinculin, play a pivotal role in facilitating mechanochemical feedback. Immunoblotting analysis of related proteins was thus carried out. Results showed elevated level of both vinculin and p-FAK of C2C12, NIH-3T3 and chondrocytes cell lines on hydrogel regions with corresponding tissue-mimicking viscoelasticity (**HAG-2/3/4**) (Fig. 5a, i-iii), but not the substrate with the highest elasticity, indicating viscosity is another contribution factor for cell-ECM interaction. Similarly, immunofluorescent analysis showed C2C12 (Fig. 5f, 5g), NIH-3T3 and chondrocytes showed higher mean fluorescence intensity (MFI) of both p-FAK and vinculin (Supplementary Fig. 13-15) and larger mean area (31 µm^2^) of FAs (Fig. 5h, i) on corresponding **HAG-2** viscoelastic region, compared to those with gradient viscoelasticity on both sides (**HAG-1** and **HAG-3**), indicating more stable focal adhesion formation within cells.

**Fig. 5:**
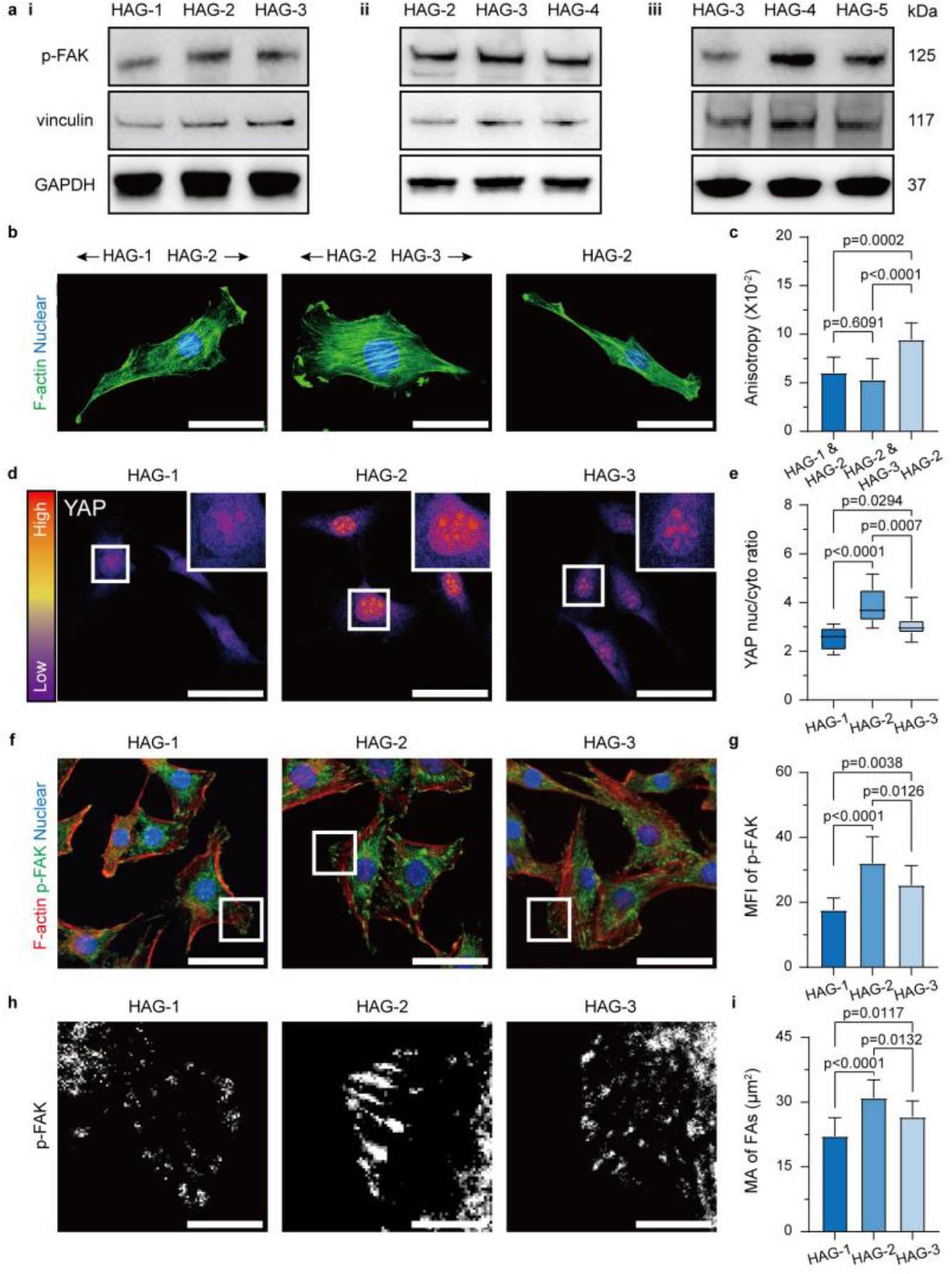
Tissue-like hydrogel increase the mechano-transduction and facilitate FAs formation. a) Western blot of i) C2C12 on **HAG-1, HAG-2** and **HAG-3**, ii) NIH-3T3 on **HAG-2, HAG-3** and **HAG-4**, and iii) chondrocytes on **HAG-3, HAG-4** and **HAG-5**, respectively. b) Immunofluorescence images of F-actin (Green) in C2C12 on **HAG-1∼HAG-2, HAG-2∼HAG-3** and **HAG-2**. Scale bar = 15 µm. c) Quantitative analysis of anisotropy in C2C12 on **HAG-1∼HAG-2, HAG-2∼HAG-3** and **HAG-2**. Error bars, mean ± s.e.m. Ordinary one-way ANOVA was used for analysis of the data; n = 12 cells. d) Immunofluorescence images of YAP in C2C12 on **HAG-1, HAG-2** and **HAG-3**. Scale bar = 30 µm. e) Quantitative analysis of YAP nuclear localization (intensity ratio between the nucleus and cytoplasm) in C2C12 on **HAG-1, HAG-2** and **HAG-3**. Error bars, mean ± s.e.m. Ordinary one-way ANOVA was used for analysis of the data; n = 15 cells. f) Immunofluorescence images of p-FAK (Green) and F-actin (Red) in C2C12 on **HAG-1, HAG-2** and **HAG-3**. Scale bar = 30 µm. g) Quantitative analysis of p-FAK mean fluorescence intensity (MFI) in C2C12 on **HAG-1, HAG-2** and **HAG-3**. Error bars, mean ± s.e.m. Ordinary one-way ANOVA was used for analysis of the data; n = 15 cells. h) Enlarged image of immunofluorescence images of p-FAK in g). i) Quantitative analysis of the mean area (MA) of FAs in C2C12 on **HAG-1, HAG-2** and **HAG-3**. Scale bar = 5 µm. Error bars, mean ± s.e.m. Ordinary one-way ANOVA was used for analysis of the data; n = 15 cells.

Interestingly, when analyzing cells at the boundary of the two distinct viscoelastic boundaries, we observed obvious anisotropy of cell stress fibers comparing to cells on homogenous substrates. The study of stress fiber anisotropy indicates heterogeneous force application when migrating across different viscoelasticity^40,41^. Indeed, the viscoelastic gradient would make a more orderly arrangement of stress fibers (Fig. 5b, 5c), which further explained the directed migration behavior of C2C12, NIH-3T3 and chondrocytes cell lines on certain gradient viscoelastic hydrogels.

Yes-associated protein (YAP), a transcriptional regulator and Hippo pathway effector protein, plays a critical role in integrating different mechanical and biochemical cues epigenetically^42^. The nuclear localization and activation of YAP are related to the increase of stress fiber assembly and FAs formation, which facilitate adhesion turnover and cell migration^43,44^. The endogenous YAP of above three cell lines on different viscoelastic hydrogels were stained to examine the viscoelasticity-induced nuclear translocation. Obviously, C2C12 displayed much more nuclear location of YAP on **HAG-2** hydrogel with muscular viscoelasticity other than **HAG-1** and **HAG-3** (Fig. 5d, 5e). YAP immunofluorescence staining image of NIH-3T3 (Supplementary Fig. 13) and chondrocytes (Supplementary Fig. 14) gave similar results, as nuclear translocation of YAP was more efficient in cells incubated on **HAG-3** and **HAG-4** hydrogels with corresponding physiological viscoelasticity, respectively. The above results indicated an enhanced YAP mediated pathway during the viscoelastic responses.

### Programmed cell migration and separation of multi-cellular systems on tissue ECM-mimic viscoelastic gradient hydrogels

Our results showed interesting migration behavior that cell traction and migration is not only rely on stiffness of the substrate. Viscosity alternation enabled physiological viscoelastic mimicking of native tissue in our system. We then wonder if through such strategy, we might spatially control cell location via such differentiated cellular behaviors (Fig. 6a). To validate this, three viscoelastic gradient **HAG** hydrogels, **HAG-a** (with viscoelasticity gradient between **HAG-2** and **HAG-3**), **HAG-b** (with viscoelasticity gradient between **HAG-3** and **HAG-4**) and **HAG-c** (with viscoelasticity gradient between **HAG-2** and **HAG-4**), were fabricated. The **HAG-a/b/c** hydrogels were divided into six areas (I-VI, Fig. 6b-6e). To distinguish different types of cells, NIH3T3 were labeled with GFP, while C2C12 were labeled with mCherry, no label was applied to chondrocytes. Cells were mixed and seeded randomly at the same time (Fig. 6b and Supplementary Fig. 16). The percentages of incubated cells on each divided area were then analyzed through 72 hours after cell plating.

**Fig. 6:**
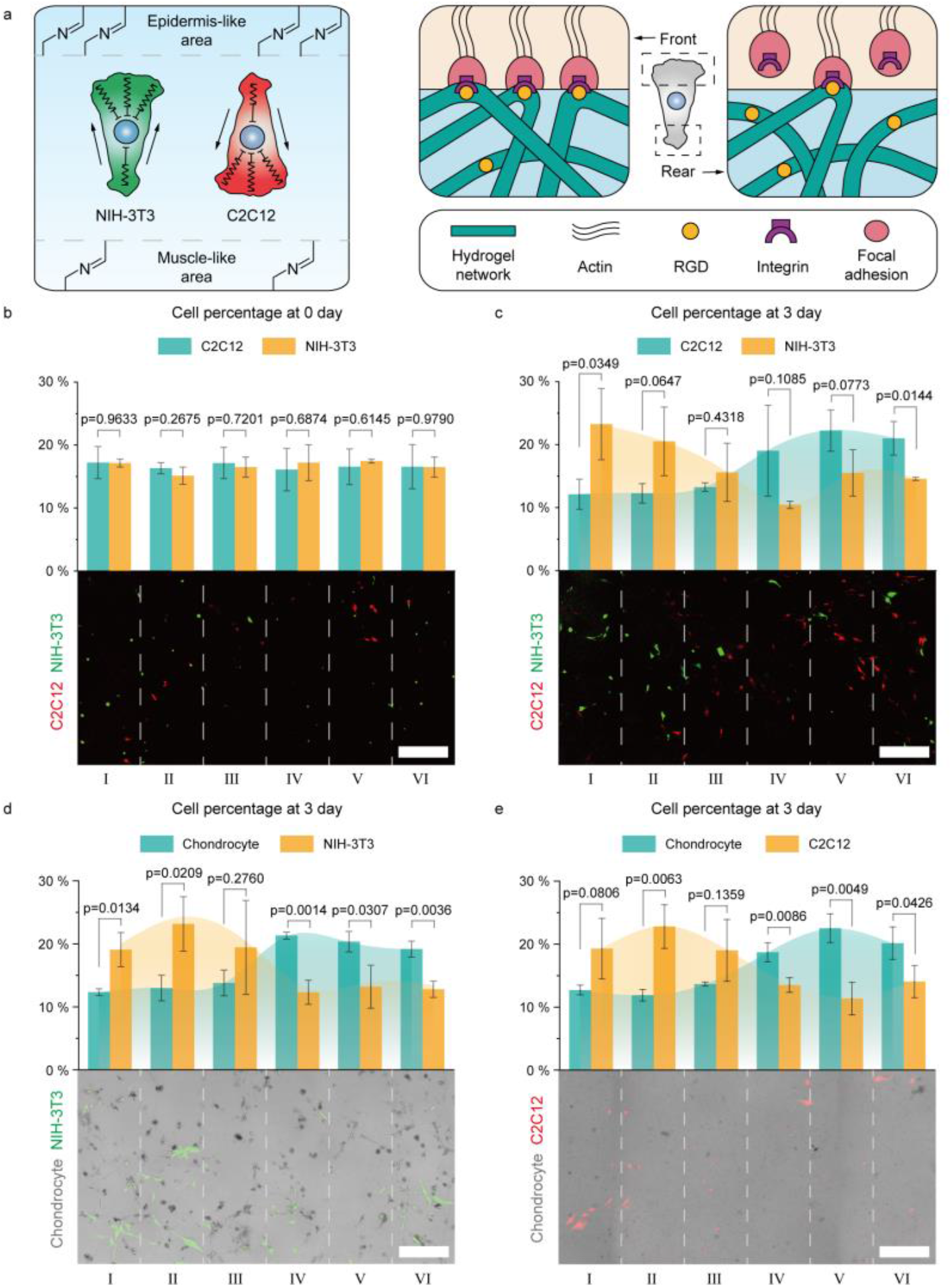
Multiple tissue-like gradient hydrogel mediated mixed cell separation. a) Diagram of mixed C2C12 and NIH-3T3 separated by **HAG-a** and the interactions between cell and matrix at cell front and rear. b) Fluorescence image and analysis of cell percentage (number of A cells in the area/total number of A cells in the figure) of C2C12 (red) and NIH-3T3 (green) seeded on **HAG-a** at Day 0. Error bars, mean ± s.e.m. Analysed by the Multiple t tests – one per row. Scale bar = 100 µm. c) Fluorescence image and analysis of cell percentage of C2C12 (red) and NIH-3T3 (green) seeded on **HAG-a** at Day 3. Error bars, mean ± s.e.m. Analysed by the Multiple t tests – one per row. Scale bar = 100 µm. d) Fluorescence image and analysis of cell percentage of chondrocyte (grey) and NIH-3T3 (green) seeded on **HAG-b** at Day 3. Error bars, mean ± s.e.m. Analysed by the Multiple t tests – one per row. Scale bar = 100 µm. e) Fluorescence image and analysis of cell percentage of chondrocyte (grey) and C2C12 (red) seeded on **HAG-c** at Day 3. Error bars, mean ± s.e.m. Analysed by the Multiple t tests – one per row. Scale bar = 100 µm.

As illustrated by Fig. 6c-6e, after 3 days of cultivation and migration, three cell lines showed different distributions on substrate with viscoelastic gradients. In specific, the percentages of C2C12 exhibited a gradual increase towards the muscular viscoelastic region in **HAG-a** (Region I-III, Fig. 6c). NIH-3T3, on the other hand, migrates to the region with the viscoelastic that mimics epidermal tissue (Region IV-VI, Fig. 6c). Similar results were also obtained in **HAG-b/c**, as chondrocytes migrate towards cartilage viscoelasticity gradient (Region IV-VI, Fig. 6d-6e). Likewise, no cell sorting performances were observed when above cell pairs were cultured on homogenous hydrogels (**HAG-2/3/4**, Supplementary Fig. 17), respectively. All of the results proved that creating nonlinear hydrogels with viscoelastic gradients indeed helps differentiated migration behaviors.

## Discussion

In this study, combined with a dual component engineering approach, we fabricated a series of tissue-mimetic hydrogels with precise viscoelasticity gradients. With such an engineering model, we showed that the viscosity and elasticity of the ECM property both influence cell mechano-sensation. In our case, C2C12, NIH-3T3, and chondrocytes show maximum cell spreading at respective optimized viscoelastic regions of the hydrogel. In agreement with this, cells can be separated during migration driven by asymmetric traction of cells spreading on the gradient hydrogels. Further immunostaining and immunoblotting experiments indicate a FAK-YAP-dependent phosphorylate pathway is also involved in cell mechano-sensation of both ECM viscosity and elasticity.

Mimicking the property of nonlinear and highly dynamic behavior of the local mechanic properties has long been a challenging issue in tissue engineering. In this research, we combined properties with two different types of polymers while maintaining the cell-ECM linker site constant. This material fabrication logic allows an ECM-mimicking, systematically adjustable viscoelastic properties with high spatial-temporal control through dynamic covalent chemical bonds. In addition, via the engineering approach we applied, it is possible to further mimic the non-linear mechanical properties in a well-controlled manner. The copolymer system and engineering system shed light on the systematic analysis of cell-ECM connection and mechano-sensation, especially the viscosity of the native environment. Indeed, our system is promising to evolve into 3D cultured system and further develop to next generation ECM mimicking for more complex tissue engineering.

## Supporting information

Supporting Information for Manuscript

## Supporting Information

A more detailed description of all materials and methods is given in the SI Appendix.

## Statistical analysis

Statistical analyses were performed with GraphPad Prism 8 software, and P values were determined by one-way ANOVA or Student’s t-test. All numerical results are presented as the mean ± standard deviation (SD). A p-value<0.05 was considered to indicate statistical significance. Each experiment used three independent **HAG**.

## Data Availability

All study data are included in the article and/or SI Appendix.

## Acknowledgments

This work is supported by the National Key R&D Program of China (2023YFF0722600), NSFC (22122803, 22378121, 22105070, 32350018), the Fundamental Research Funds for the Central Universities (222201717003). JZ acknowledges Shanghai Natural Science Foundation Project (23ZR1479500, 23JC1401700). ZL acknowledges Shanghai Sailing Program (20YF1410300).

## Author Contributions

Z.L., Y.G., H.T., and J.Z. designed the research; Z.L. and J.Z. designed the hydrogel engineering approach; H.Y., M.L. and Y.S. synthesized the hydrogel; H.Y., M.L. and L.C. performed all biological experiments; Z.L., H.Y., Y.G., and J.Z. analyzed data; Y.G., J.Z. and H.T. wrote the paper.

