## Supporting Information for Manuscript for "Engineering tissue-mimetic hydrogels with continuous viscoelastic gradient for programmed cell migration"

^1^Key Laboratory for Advanced Materials and Joint International Research Laboratory of Precision Chemistry and Molecular Engineering, Feringa Nobel Prize Scientist Joint Research Center, Frontiers Science Center for Materiobiology and Dynamic Chemistry, Institute of Fine Chemicals, School of Chemistry and Molecular Engineering, East China University of Science and Technology, Shanghai 200237, China

^2^Y. Shang, Y. Ge

^2^Interdisciplinary Research Center on Biology and Chemistry, Shanghai Institute of Organic Chemistry, Chinese Academy of Sciences, Shanghai 201210, China

^3^ Y. Shang current address: School of Chemistry and Molecular Engineering, East China Normal University, Shanghai 200241, China

Table of Contents

1. **Experimental information for material**	1
2. **Experimental information for cell experiment**	12
3. **References**	18

**1. Experimental information for material**

**1.1 Preparation of Decellularized matrix**

According to previous reported method^1,2,3^, 250–350 g rats were used as a source of muscle and epidermis for decellularisation. Rats were killed by CO_2_ inhalation and death confirmed by onset of rigor mortis. The mouse latissimus dorsi muscles obtained from 3 rats were treated with 0.25 % SDS for 72 hours at 4 °C and washed in de-ionized water for 48 hours to obtain decellularized muscle matrix and the dorsi epidermis obtained from 3 rats were treated with trysin and triton X-100 to obtain decellularized epidermis matrix. Cartilage obtained from rabbit ear were cut into pieces and maintained at − 80 °C for three days, and then the tissue specimen was snap frozen and thawed in liquid nitrogen four times at 2-min intervals. After that, tissues were treated with 5% SDS for 8 h to obtain decellularized cartilage matrix.

**1.2 Preparation of oxi-HA**

Oxi-HA was synthesized according to the previously reported method^4,5^. Briefly, 1 g HA (80-150 kDa) as dissolved in 100 mL deionized water at room temperature, then 2.24 g NaIO_4_ was solved in 15 mL deionized water and added slowly to the HA solution. The mixture was stirred for 24 h in a dark environment at room temperature and then quenched with 1 mL ethylene glycol for a half hour. The resulting product was dialyzed against deionized water for 7 days by using a semipermeable membrane (with a MWCO of 14 kDa). Finally, the dialyzed solution was lyophilized, yielding a white fluffy product.

**1.3 Determination of the oxidation degree**

The oxidation degree (OD) was quantified by measuring the number of aldehyde functional groups in oxi-HA using t-BC and TNBS according to the previous method^4,5^. Briefly, 0.5 mL of oxi-HA (0.6% w/v) and 0.5 mL of t-BC (30 mM) in 1% aqueous trichloroacetic acid were well mixed and reacted at room temperature for 24 h. Then, 0.1 M of borate buffer was used to dilute the reaction mixture (1:100), 500 μL 0.1% (v/v) of TNBS and 500 μL diluted reaction mixture was mixed and reacted for 3 hours at 37°C. Then, the solution was diluted with hydrochloric acid (0.5N) and the absorbance of the solution was measured at 330 nm. A standard calibration curve was obtained with t-BC solutions ranging from 0 M up to 30 mM to determine the amount of unreacted t-BC and to calculate the OD.

y=0.3266x+0.01776, R^2^=0.9980	(1)

where y is the absorbance, and x is the t-BC concentration (g/L)

$\mathrm{OD}\left( \% \right)=\left( \frac{n_{0}-n_{1}}{n_{0}} \right)$×100%	(2)

Where n_0_ and n_1_ are the initial and final number of moles of t-BC, respectively. As can been seen in Supplementary Fig. 1, the OD of oxi-HA was 84%.

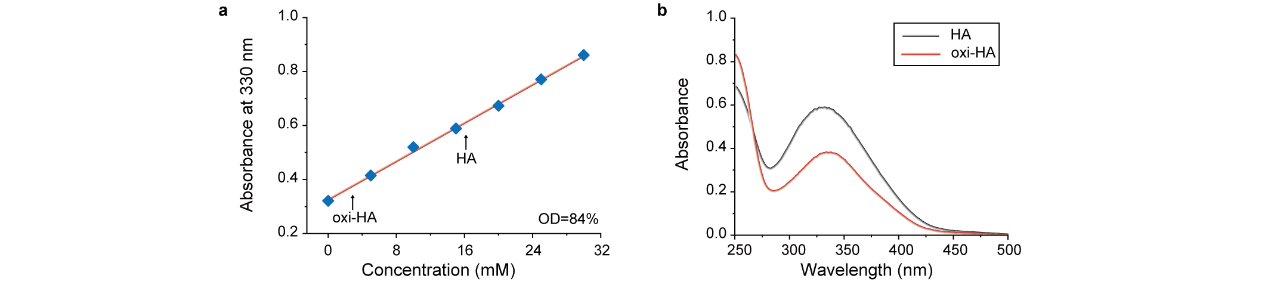

**Supplementary Fig. 1** a) The standard calibration curve of t-BC (0-30 mM) and TNBS (0.1 % v/v). b) The absorbance of HA and oxi-HA react with TBBS, respectively.

**1.4 Preparation of the HAG hydrogel**

Different weight of oxi-HA was dissolved overnight in PBS (pH=7.4) and different weight of gelatin was dissolved in PBS at 37°C to obtain different concentration of solution. Then, the oxi-HA and gelatin solutions were mixed homogeneously and put in a dark environment for 1 day at room temperature to form the C=N bond.

**1.5 Characterization of functional groups**

The functional groups of oxi-HA and HAG hydrogel with different component were identified by Fourier transform infrared (FTIR) spectroscopy using a Nicolet 6700 (Thermo Scientific, Madison, USA) instrument equipped with an attenuated total reflectance (ATR) accessory and KBr crystal. The spectra were obtained in the range of 800-2000 cm^-1^. The samples were previously lyophilized for 24 h and immediately analyzed.

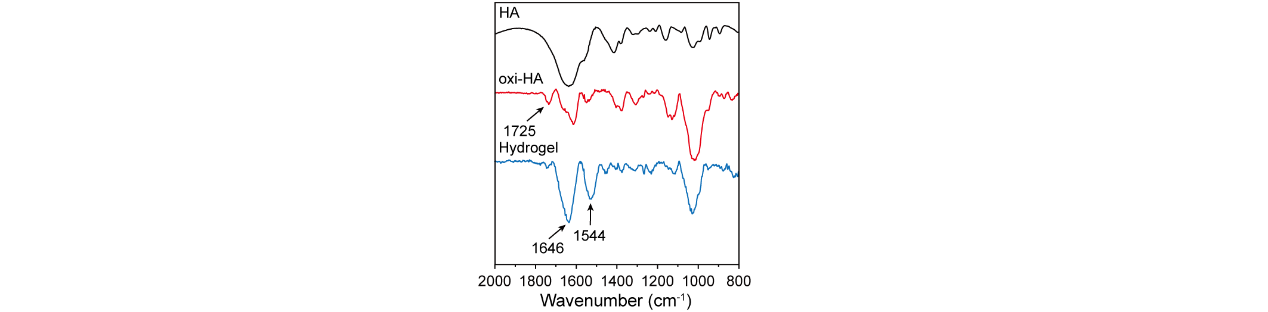

**Supplementary Fig. 2** FT-IR spectrum of HA, oxi-HA and hydrogel (HAG-1).

**1.6 Morphology**

The HAG samples were lyophilized and kept in a desiccator to obtain the images. The morphology of the HAG hydrogels was obtained by scanning electron microscopy (SEM) (S-4800, Hitachi, Japan) using a voltage of 15 kV. The average sizes of the pores were determined by measuring 21 individual pores from three different images using ImageJ.

**1.7 Characterization of HAG viscoelasticity by rheometer**

Rheological characterizations were performed using a Discovery HR-2 (TA, USA). Samples were placed between the bottom rheometer plate and a 20 mm flat plate (1 mm thick). Experiments were performed at 25 °C. The oscillation frequency and oscillation time were performed at 0.1% strain. The step stress relaxation was performed at 0.08% strain.

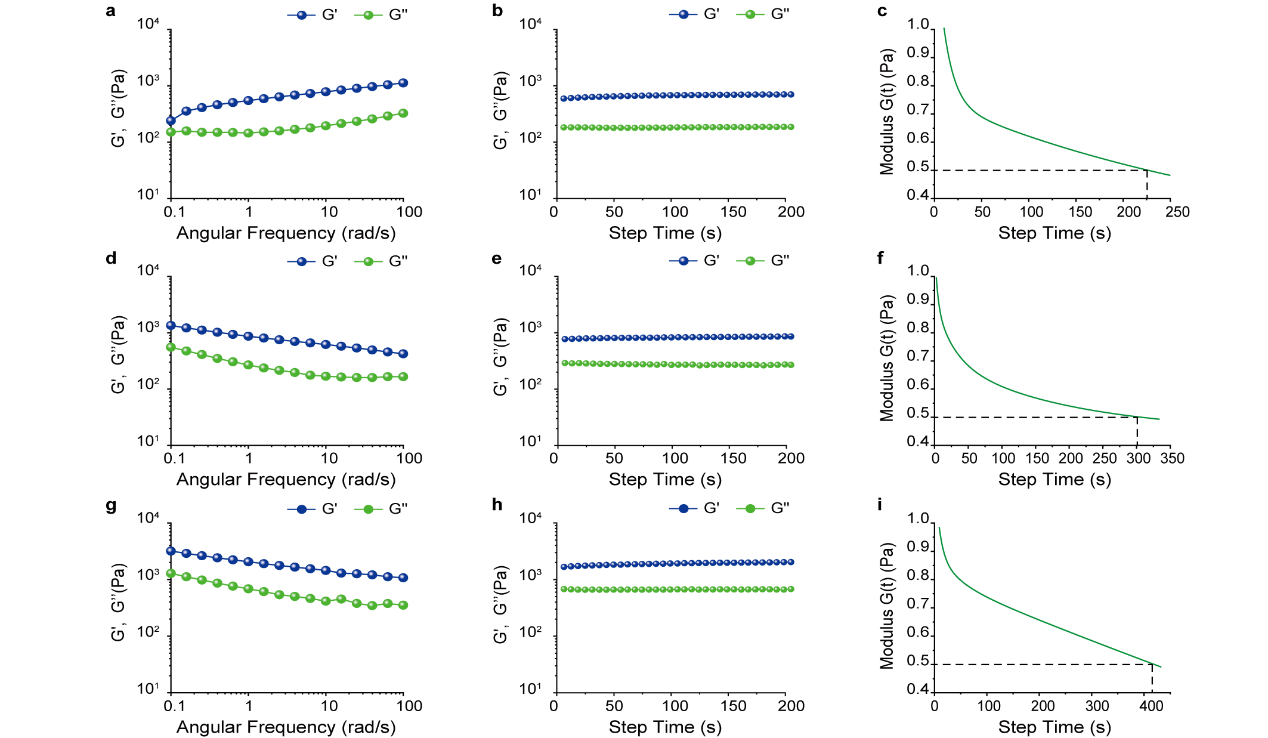

**Supplementary Fig. 3** The viscoelastic properties of decellularized muscle, where a) refer to the frequency sweep (0.1% strain), b) refer to the time sweep (0.1% strain) and c) refer to the stress relaxation curves (0.08% strain). The viscoelastic properties of decellularized epidermis, where d) refer to the frequency sweep (0.1% strain), e) refer to the time sweep (0.1% strain) and f) refer to the stress relaxation curves (0.08% strain). The viscoelastic properties of decellularized cartilage, where g) refer to the frequency sweep (0.1% strain), h) refer to the time sweep (0.1% strain) and i) refer to the stress relaxation curves (0.08% strain).

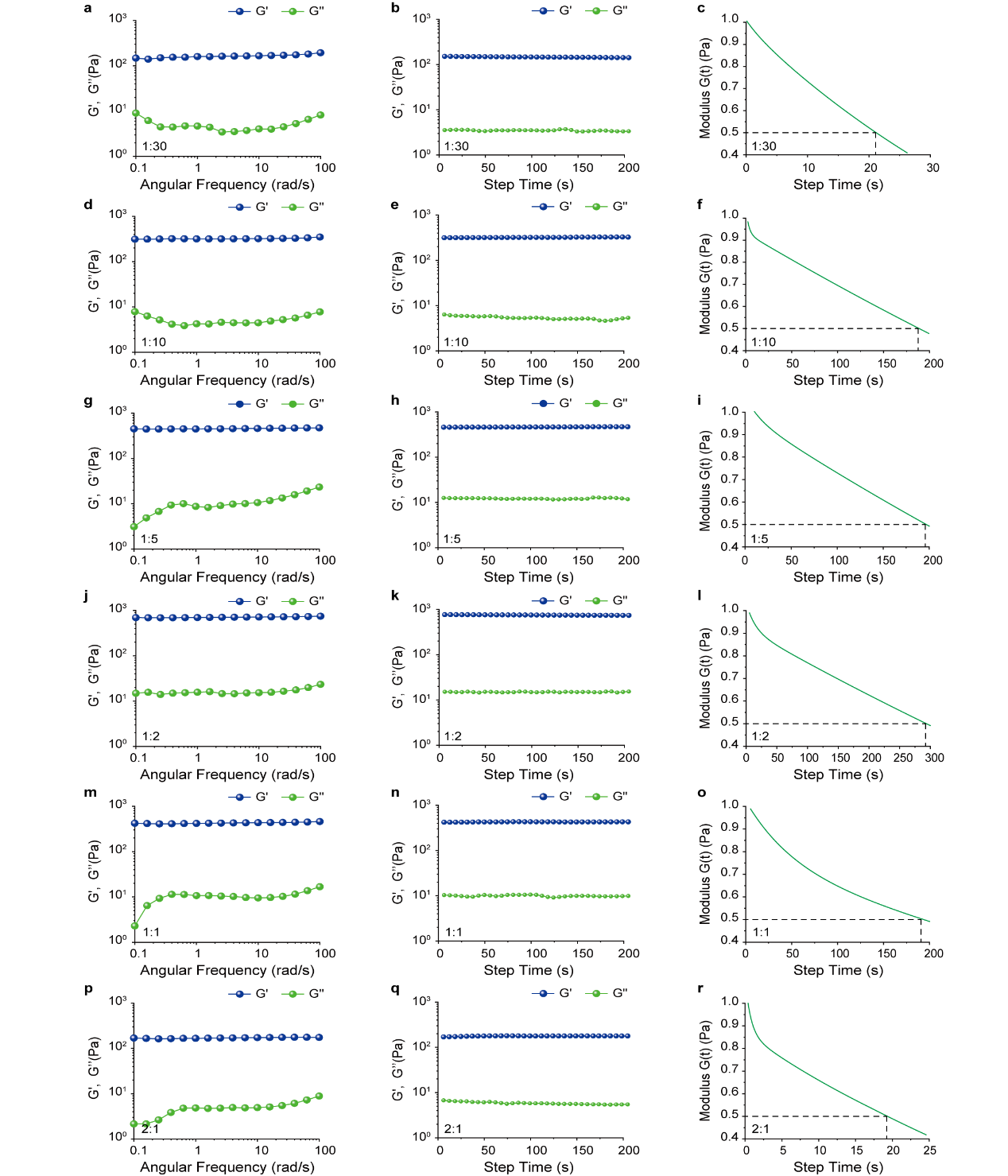

**Supplementary Fig. 4** The viscoelastic properties of 10 % HAG, which mass ratio between oxi-HA and Gelatin was 1:30, where a) refer to the frequency sweep (0.1% strain), b) refer to the time sweep (0.1% strain) and c) refer to the stress relaxation curves (0.08% strain). The viscoelastic properties of 10 % HAG, which mass ratio between oxi-HA and Gelatin was 1:10, where d) refer to the frequency sweep (0.1% strain), e) refer to the time sweep (0.1% strain) and f) refer to the stress relaxation curves (0.08% strain). The viscoelastic properties of 10 % HAG, which mass ratio between oxi-HA and Gelatin was 1:5, where g) refer to the frequency sweep (0.1% strain), h) refer to the time sweep (0.1% strain) and i) refer to the stress relaxation curves (0.08% strain). The viscoelastic properties of 10 % HAG, which mass ratio between oxi-HA and Gelatin was 1:2, where j) refer to the frequency sweep (0.1% strain), k) refer to the time sweep (0.1% strain) and l) refer to the stress relaxation curves (0.08% strain). The viscoelastic properties of 10 % HAG, which mass ratio between oxi-HA and Gelatin was 1:1, where m) refer to the frequency sweep (0.1% strain), n) refer to the time sweep (0.1% strain) and o) refer to the stress relaxation curves (0.08% strain). The viscoelastic properties of 10 % HAG, which mass ratio between oxi-HA and Gelatin was 2:1, where p) refer to the frequency sweep (0.1% strain), q) refer to the time sweep (0.1% strain) and r) refer to the stress relaxation curves (0.08% strain).

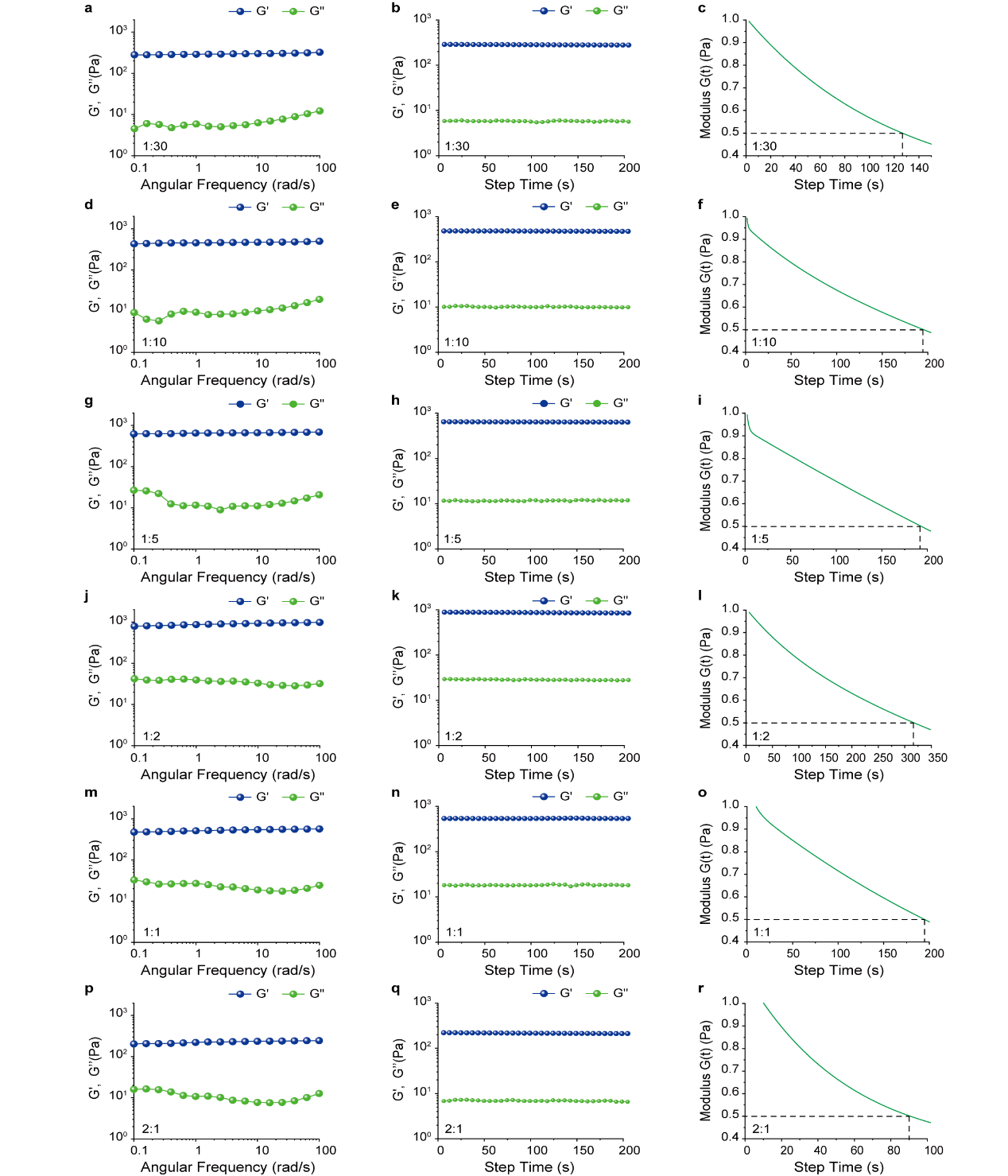

**Supplementary Fig. 5** The viscoelastic properties of 13 % HAG, which mass ratio between oxi-HA and Gelatin was 1:30, where a) refer to the frequency sweep (0.1% strain), b) refer to the time sweep (0.1% strain) and c) refer to the stress relaxation curves (0.08% strain). The viscoelastic properties of 13 % HAG, which mass ratio between oxi-HA and Gelatin was 1:10, where d) refer to the frequency sweep (0.1% strain), e) refer to the time sweep (0.1% strain) and f) refer to the stress relaxation curves (0.08% strain). The viscoelastic properties of 13 % HAG, which mass ratio between oxi-HA and Gelatin was 1:5, where g) refer to the frequency sweep (0.1% strain), h) refer to the time sweep (0.1% strain) and i) refer to the stress relaxation curves (0.08% strain). The viscoelastic properties of 13 % HAG, which mass ratio between oxi-HA and Gelatin was 1:2, where j) refer to the frequency sweep (0.1% strain), k) refer to the time sweep (0.1% strain) and l) refer to the stress relaxation curves (0.08% strain). The viscoelastic properties of 13 % HAG, which mass ratio between oxi-HA and Gelatin was 1:1, where m) refer to the frequency sweep (0.1% strain), n) refer to the time sweep (0.1% strain) and o) refer to the stress relaxation curves (0.08% strain). The viscoelastic properties of 13 % HAG, which mass ratio between oxi-HA and Gelatin was 2:1, where p) refer to the frequency sweep (0.1% strain), q) refer to the time sweep (0.1% strain) and r) refer to the stress relaxation curves (0.08% strain).

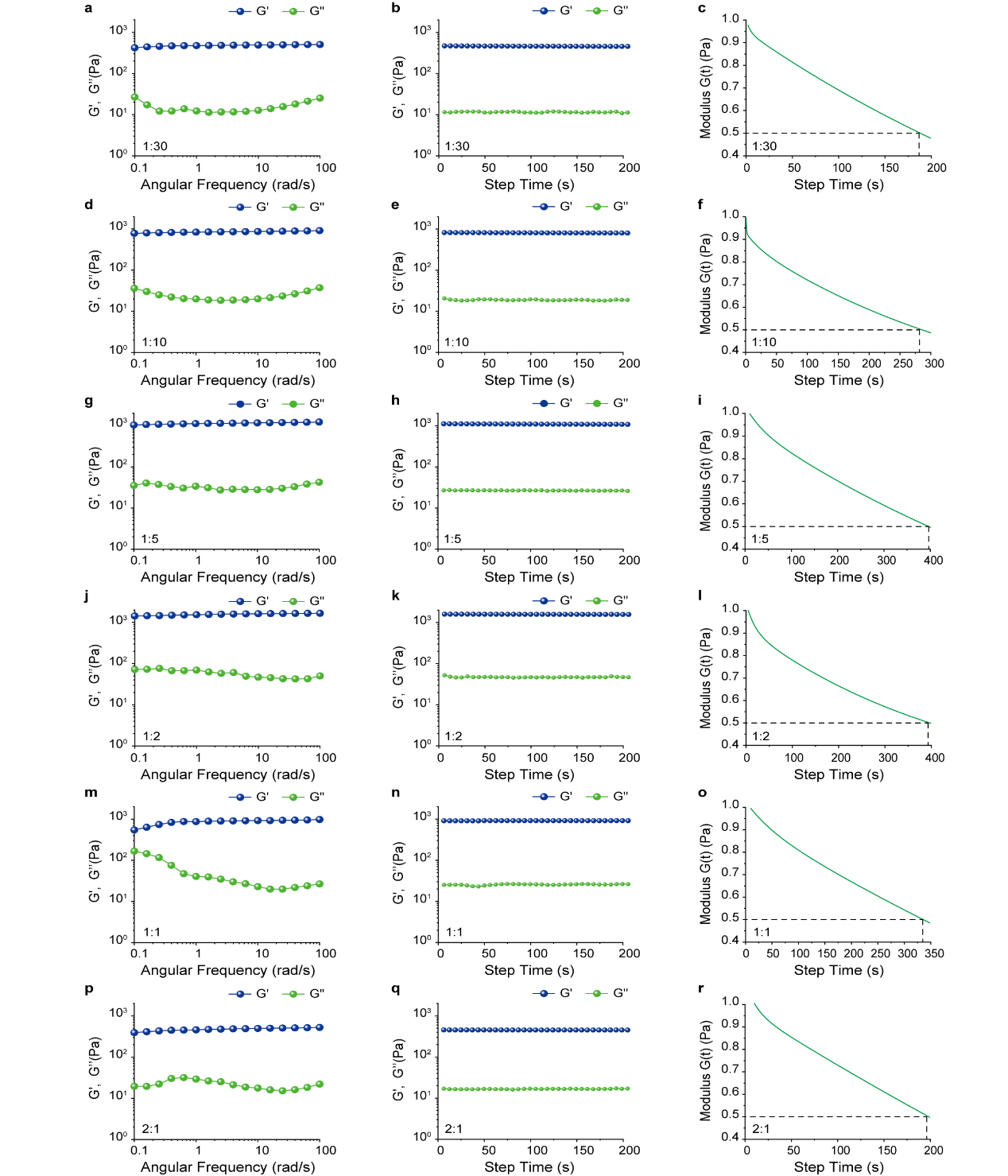

**Supplementary Fig. 6** The viscoelastic properties of 15 % HAG, which mass ratio between oxi-HA and Gelatin was 1:30, where a) refer to the frequency sweep (0.1% strain), b) refer to the time sweep (0.1% strain) and c) refer to the stress relaxation curves (0.08% strain). The viscoelastic properties of 15 % HAG, which mass ratio between oxi-HA and Gelatin was 1:10, where d) refer to the frequency sweep (0.1% strain), e) refer to the time sweep (0.1% strain) and f) refer to the stress relaxation curves (0.08% strain). The viscoelastic properties of 15 % HAG, which mass ratio between oxi-HA and Gelatin was 1:5, where g) refer to the frequency sweep (0.1% strain), h) refer to the time sweep (0.1% strain) and i) refer to the stress relaxation curves (0.08% strain). The viscoelastic properties of 15 % HAG, which mass ratio between oxi-HA and Gelatin was 1:2, where j) refer to the frequency sweep (0.1% strain), k) refer to the time sweep (0.1% strain) and l) refer to the stress relaxation curves (0.08% strain). The viscoelastic properties of 15 % HAG, which mass ratio between oxi-HA and Gelatin was 1:1, where m) refer to the frequency sweep (0.1% strain), n) refer to the time sweep (0.1% strain) and o) refer to the stress relaxation curves (0.08% strain). The viscoelastic properties of 15 % HAG, which mass ratio between oxi-HA and Gelatin was 2:1, where p) refer to the frequency sweep (0.1% strain), q) refer to the time sweep (0.1% strain) and r) refer to the stress relaxation curves (0.08% strain).
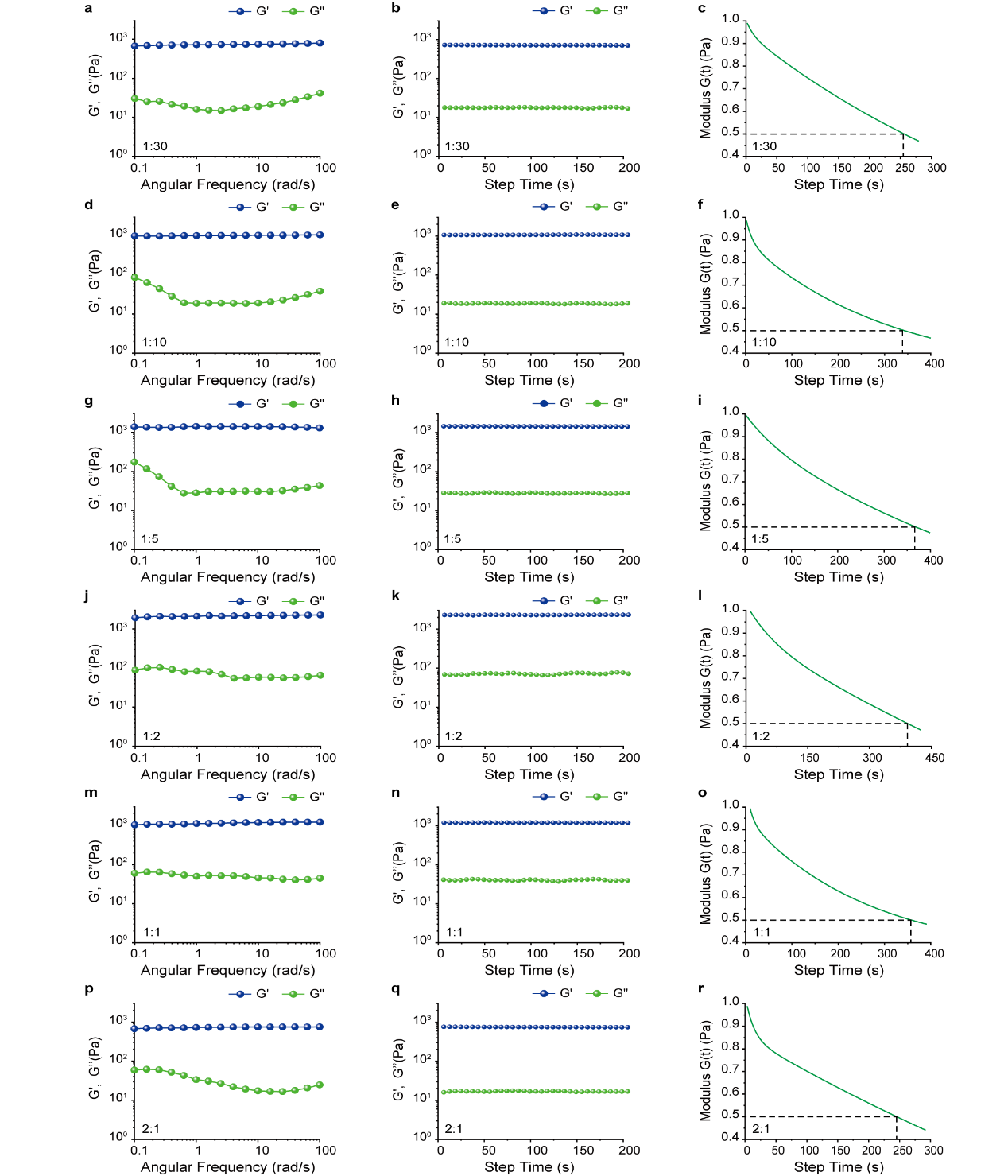
**Supplementary Fig. 7** The viscoelastic properties of 17 % HAG, which mass ratio between oxi-HA and Gelatin was 1:30, where a) refer to the frequency sweep (0.1% strain), b) refer to the time sweep (0.1% strain) and c) refer to the stress relaxation curves (0.08% strain). The viscoelastic properties of 17 % HAG, which mass ratio between oxi-HA and Gelatin was 1:10, where d) refer to the frequency sweep (0.1% strain), e) refer to the time sweep (0.1% strain) and f) refer to the stress relaxation curves (0.08% strain). The viscoelastic properties of 17 % HAG, which mass ratio between oxi-HA and Gelatin was 1:5, where g) refer to the frequency sweep (0.1% strain), h) refer to the time sweep (0.1% strain) and i) refer to the stress relaxation curves (0.08% strain). The viscoelastic properties of 17 % HAG, which mass ratio between oxi-HA and Gelatin was 1:2, where j) refer to the frequency sweep (0.1% strain), k) refer to the time sweep (0.1% strain) and l) refer to the stress relaxation curves (0.08% strain). The viscoelastic properties of 17 % HAG, which mass ratio between oxi-HA and Gelatin was 1:1, where m) refer to the frequency sweep (0.1% strain), n) refer to the time sweep (0.1% strain) and o) refer to the stress relaxation curves (0.08% strain). The viscoelastic properties of 17 % HAG, which mass ratio between oxi-HA and Gelatin was 2:1, where p) refer to the frequency sweep (0.1% strain), q) refer to the time sweep (0.1% strain) and r) refer to the stress relaxation curves (0.08% strain).

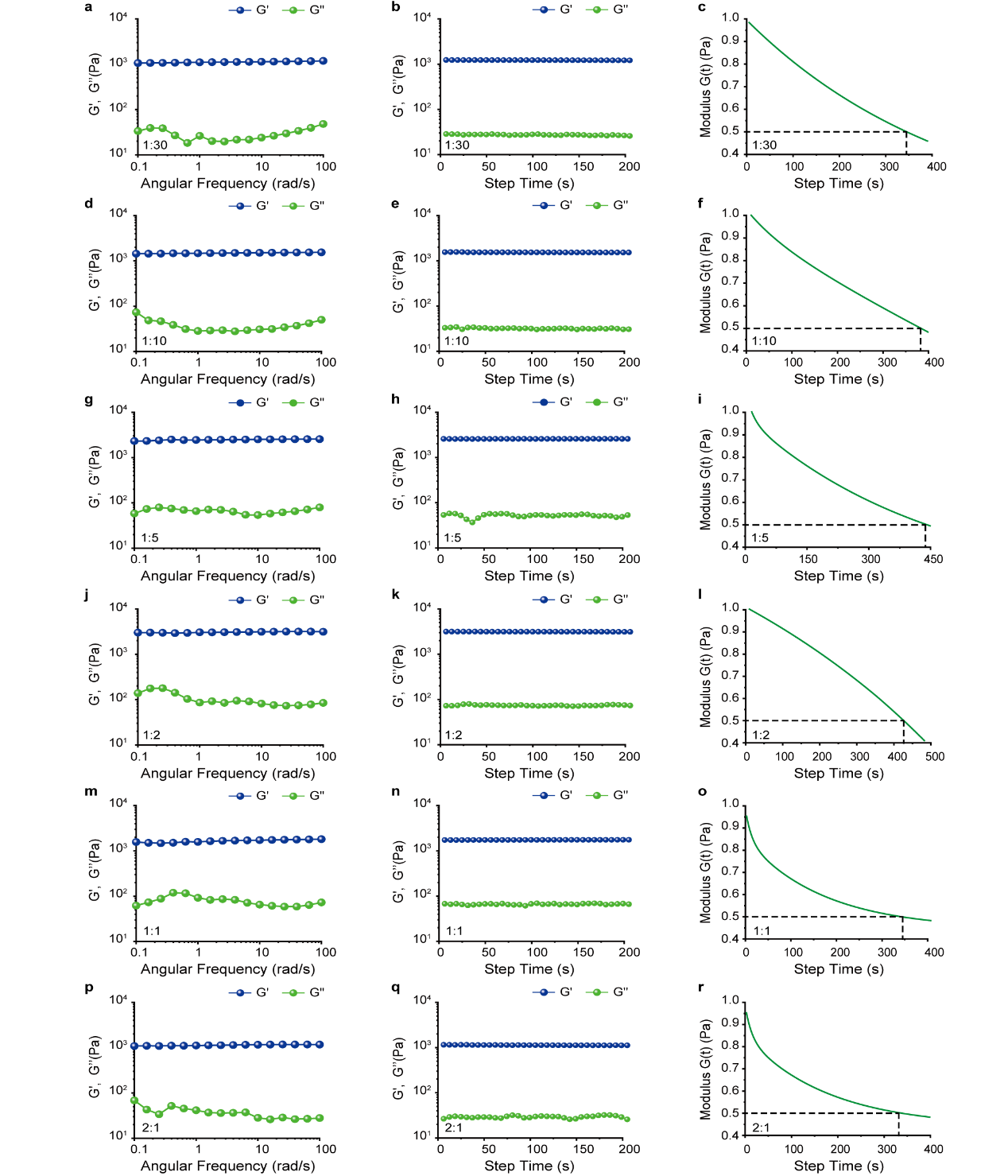
 **Supplementary Fig. 8** The viscoelastic properties of 20 % HAG, which mass ratio between oxi-HA and Gelatin was 1:30, where a) refer to the frequency sweep (0.1% strain), b) refer to the time sweep (0.1% strain) and c) refer to the stress relaxation curves (0.08% strain). The viscoelastic properties of 20 % HAG, which mass ratio between oxi-HA and Gelatin was 1:10, where d) refer to the frequency sweep (0.1% strain), e) refer to the time sweep (0.1% strain) and f) refer to the stress relaxation curves (0.08% strain). The viscoelastic properties of 20 % HAG, which mass ratio between oxi-HA and Gelatin was 1:5, where g) refer to the frequency sweep (0.1% strain), h) refer to the time sweep (0.1% strain) and i) refer to the stress relaxation curves (0.08% strain). The viscoelastic properties of 20 % HAG, which mass ratio between oxi-HA and Gelatin was 1:2, where j) refer to the frequency sweep (0.1% strain), k) refer to the time sweep (0.1% strain) and l) refer to the stress relaxation curves (0.08% strain). The viscoelastic properties of 20 % HAG, which mass ratio between oxi-HA and Gelatin was 1:1, where m) refer to the frequency sweep (0.1% strain), n) refer to the time sweep (0.1% strain) and o) refer to the stress relaxation curves (0.08% strain). The viscoelastic properties of 20 % HAG, which mass ratio between oxi-HA and Gelatin was 2:1, where p) refer to the frequency sweep (0.1% strain), q) refer to the time sweep (0.1% strain) and r) refer to the stress relaxation curves (0.08% strain).

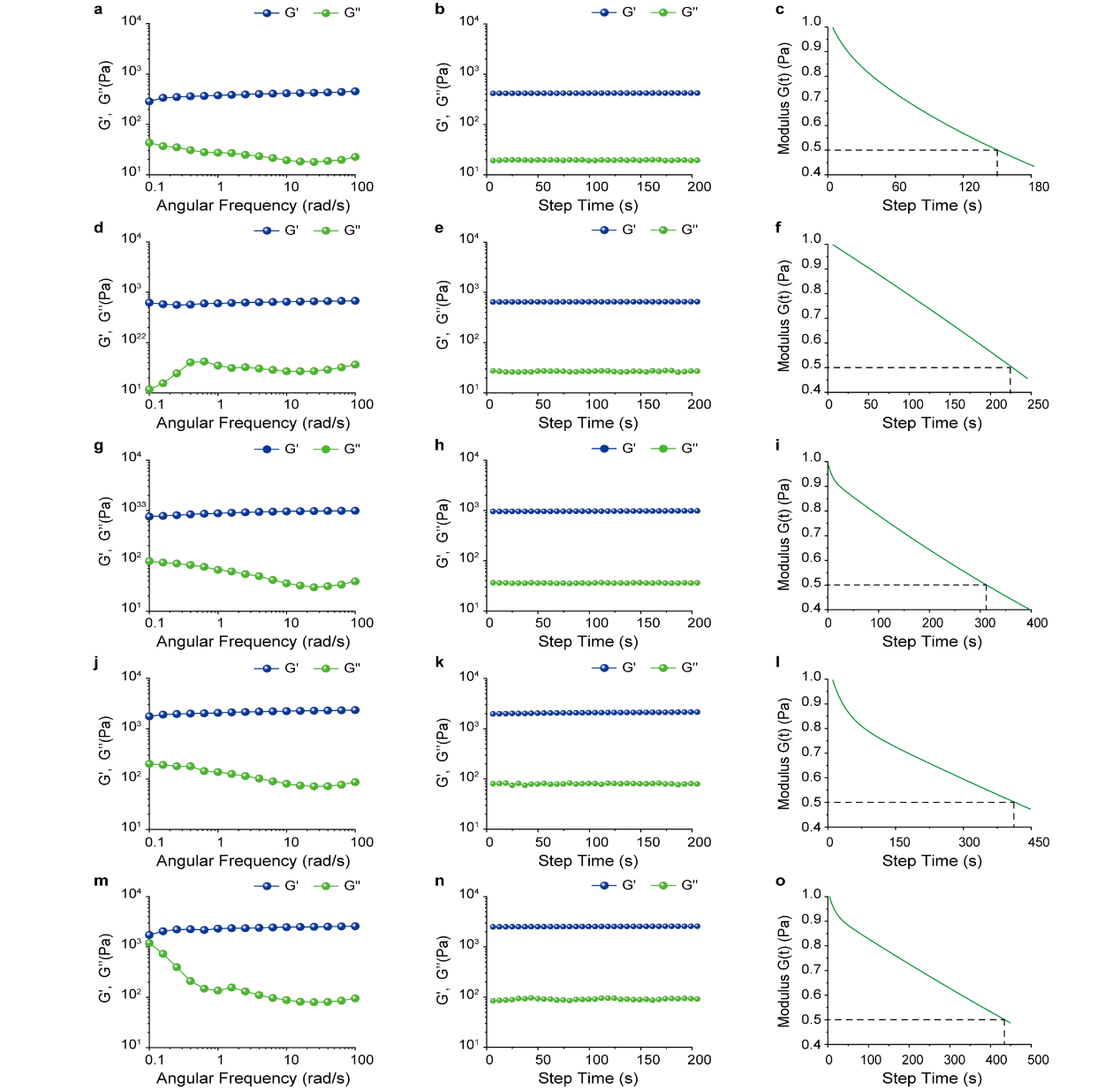
 **Supplementary Fig. 9** The viscoelastic properties of HAG-1, where a) refer to the frequency sweep (0.1% strain), b) refer to the time sweep (0.1% strain) and c) refer to the stress relaxation curves (0.08% strain). The viscoelastic properties of HAG-2, where d) refer to the frequency sweep (0.1% strain), e) refer to the time sweep (0.1% strain) and f) refer to the stress relaxation curves (0.08% strain). The viscoelastic properties of HAG-3, where g) refer to the frequency sweep (0.1% strain), h) refer to the time sweep (0.1% strain) and i) refer to the stress relaxation curves (0.08% strain). The viscoelastic properties of HAG-4, where j) refer to the frequency sweep (0.1% strain), k) refer to the time sweep (0.1% strain) and l) refer to the stress relaxation curves (0.08% strain). The viscoelastic properties of HAG-5, where m) refer to the frequency sweep (0.1% strain), n) refer to the time sweep (0.1% strain) and o) refer to the stress relaxation curves (0.08% strain).

**1.8 Characterization of HAG viscoelasticity by nano-indentation**

The characterization of viscoelastic gradient hydrogel were performed using an OP1550 interferometer. For the investigation of effective Young's modulus, the indentation cycle followed by a hold period of 1 s at maximum load and the unloading part was 2 s. For the measurement of stress relaxation time, the hold period was 10-150 s at maximum load according to different HAG and the unloading part was 2s. Experiments were performed at 25 °C.

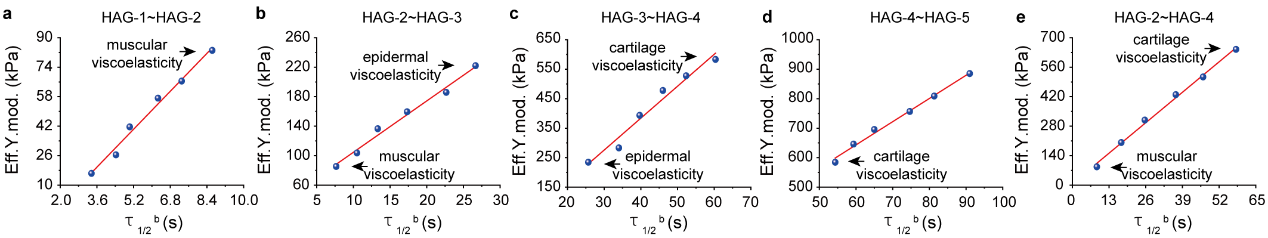

**Supplementary Fig. 10** Linear fitting results of effective Young's modulus and stress relaxation time of gradient viscoelastic hydrogel. a) The linear equation of HAG-1~HAG-2 was y=12865x-26570, Adj.R^2^=0.98393, b) The linear equation of HAG-2~HAG-3 was y=6965x+34908, Adj.R^2^=98392, c) The linear equation of HAG-3~HAG-4 was y=10826x-48803, Adj.R^2^=0.96227, d) The linear equation of HAG-4~HAG-5 was y=7865x+171476, Adj.R^2^=0.99069 and e) The linear equation of HAG-2~HAG-4 was y=11116x+7545, Adj.R^2^=0.99469.

**2. Experimental information for cell experiment**

**2.1 Cell culture**

C2C12, C2C12- RGP (red fluorescent protein), NIH-3T3, NIH-3T3-GFP (Green fluorescent protein) and chondrocytes were purchased from Shanghai Jinyuan Biotechnology Co., Ltd and cultured in high-glucose Dulbecco’s modifed Eagle’s medium (DMEM) supplemented with 10% fetal bovine serum (Gibco, 10099141C) and 1% penicillin/streptomycin. The cells were cultured at 37 °C**,** 5% CO_2_ in a humidified incubator.

**2.2 Cytotoxicity measurements for the HAG**

Cytotoxicity was measured according to previous method^6,7^. The prepared HAG was immersed in DMEM at 37 °C for 72 h and the extract was collected for cytotoxicity measurements. C2C12, NIH-3T3 and chondrocytes were seeded in 98-well plates at a concentration of 40000 cells per well and cultured in DMEM (10% FBS) for 24 h at 37 °C with 5% humidified CO_2_. The culture medium was replaced by the extract or the control DMEM. At 1, 3, 5 and 7 days, cells were incubated with 10 μL of Cell Counting Kit-8 (CCK-8) reagent in 5% humidified CO_2_ for 2h at 37 °C. The absorbance at 450 nm was measured by a microplate reader (Molecular Devices, Sunnyvale, CA).

Cell viability=$\frac{A_{i}}{A_{o}}$x100%	(3)

Where A_i_ is the absorbance of cells treated with the hydrogel extraction solution and A_0_ is the absorbance of the blank control.

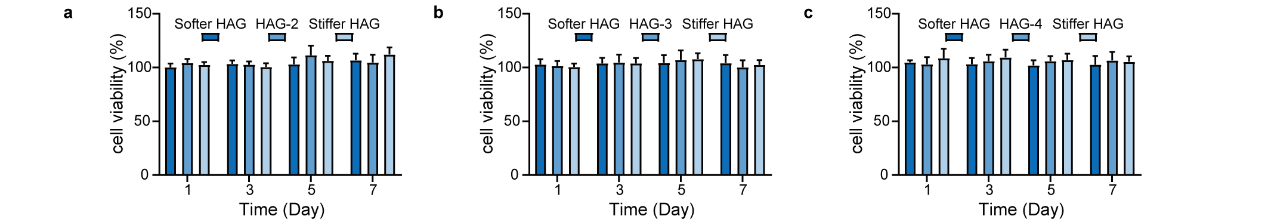

**Supplementary Fig. 11** Cell viability of a) C2C12, b) NIH-3T3 and c) chondrocyte exposed to hydrogel extract for day 1, day 3, day 5 and day 7.

**2.3 Cell migration on viscoelastic gradient substrates**

For the analysis of cells migration on gradient viscoelastic hydrogel, 20000 (C2C12 and NIH-3T3) or 30000 (chondrocytes) cells were seeded on gradient viscoelastic hydrogel and cultured to adhere for overnight. Imaging was started 12 hours after seeding and time-lapse movies were acquired for 12.5 h at 6 min intervals. For control experiments, C2C12, NIH-3T3 and chondrocytes were seeded on HAG-2, HAG-3 and HAG-4, respectively, and the time-lapse movies were also acquired at 6 min intervals for 12.5 h. After the experiment was finished, the culture was fixed and prepared for subsequent experiments, as described below. Migration tracks from individual cells were investigated for angular displacements. Mitotic, dying or crowded cells were excluded from the analysis.

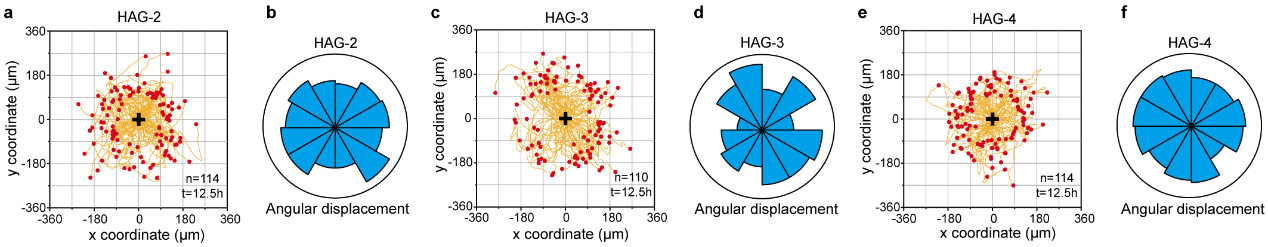

**Supplementary Fig. 12** a) Migration trajectory of C2C12 on homogeneous **HAG-2**, n = 114. b) Angular displacement of C2C12 in homogenous **HAG-2**. c) Migration trajectory of NIH-3T3 on homogenous **HAG-3**, n = 110. d) Angular displacement of NIH-3T3 in homogenous **HAG-3**. e) Migration trajectory of chondrocytes on homogenous **HAG-4**, n = 114. f) Angular displacement of chondrocytes in homogenous **HAG-4**.

**2.4 Western blotting**

Cells on hydrogels were placed on ice, rinsed twice with ice-cold phosphate-buffered saline (PBS) and scraped into a lysis buffer (A05-K1123). The lysates were vortexed, heat (90 °C) for 10 min and sonicated before separation by SDS–polyacrylamide gel electrophoresis. Then, the proteins were transferred to nitrocellulose membranes and the membranes were blocked with 5 % skimmed milk in tris-buffered saline and incubated with the indicated primary antibodies (pFAK, 44-624G, Thermo Fisher Scientific; vinculin, sc-25336, Santa Cruz Biotechnology) overnight at 4 °C, followed by fluorophore-conjugated secondary antibodies (Goat Anti-Mouse ,YFSA01; Goat Anti-Rabbit, YFSA02) for 2 h at room temperature. Finally, the membranes were scanned using a Chemiluminescence imaging system (JP-K300).

**2.5 Immunofluorescence staining**

Samples were fixed for 10 min with warm 4% paraformaldehyde, followed by permeabilization with 0.1% Triton X-100 for 10 min and blocking with 5 % skimmed milk for 30 min. Samples were incubated with the primary antibody (diluted in 5 % skimmed milk) overnight at 4 °C. Secondary antibodies were diluted in PBS and the samples were incubated with the antibody for 2 h at room temperature. The nuclei were counterstained using DAPI and filamentous actin using phalloidin (TraKine™ F-actin Staining Kit, KTC4009).

**2.6 Traction force microscopy**

To measure the tractions exerted by cells on different HAG, polystyrene fluorescent microspheres with a diameter of 200 nm were uniformly mixed in the HAG. The cells were seeded on the gels (1000 cells per plate) and grown for 24h before the experiment was conducted. For imaging the cells and microspheres, a stage-top incubator unit (37 °C/5% CO_2_) was used. Bright-field and fluorescent images of single cells were captured before and after cell detachment by the addition of 1% SDS.

Traction force microscopy was performed using the PIV and Fourier transform traction cytometry (FTTC) ImageJ plugins, called by a custom-written macro to process stacks according to previous research^[8,9]^. FTTC was performed using a Poisson ratio of 0.5 and an effective Young’s modulus value appropriate for the hydrogel being analyzed. Cell contours in phase contrast images were outlined manually and transferred to traction force maps; values outside the contours were cleared to depict only forces within the area of the cells themselves.

**2.7 Image analysis**

Images of bright field and fluorescence field were taken for the indicated conditions at X40 or X10 with a laser scanning confocal microscope (STELLARIS 8). Images were nalysed using ImageJ (64-bit) (National Institutes of Health) software. For the analysis of single cell migration trajectory, manual tracking was used to analyse the time-lapse movies. To analyse the anisotropy of F-actin, the FibriTool was used according to previous reported research ^[10]^. For the analysis of YAP nuclear localization, the mean grey value in the nucleus was divided by the corresponding mean value in the cytoplasm. For the analysis of pFAK and vinculin, background removal and thresholding were used to exclude cytoplasmic signals and the FMI was recorded.

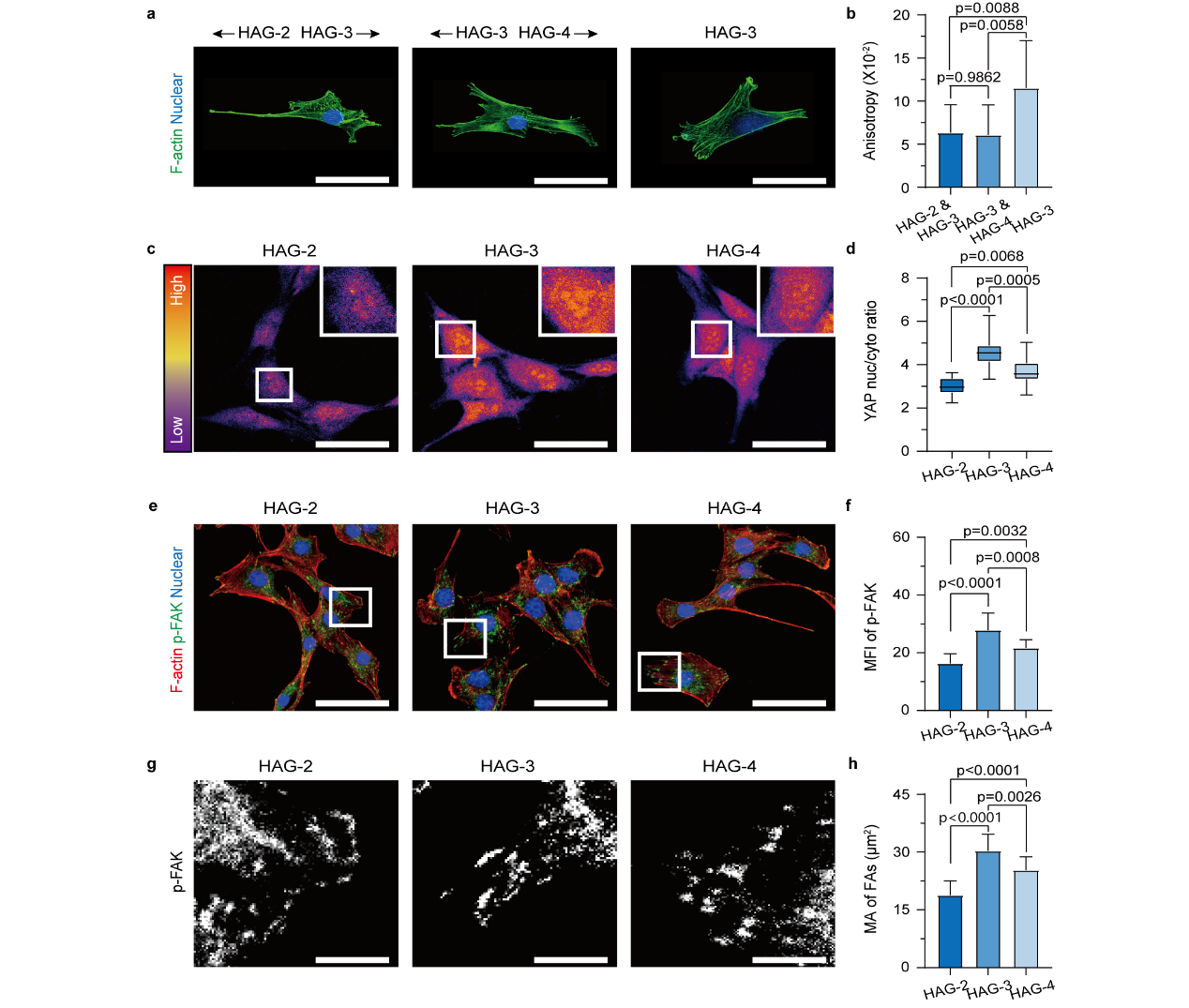

**Supplementary Fig. 13** a) Immunofluorescence images of F-actin (Green) in NIH-3T3 on HAG-2~HAG-3, HAG-3~HAG-4 and HAG-3. Scale bar=15 µm. b) Quantitative analysis of anisotropy in C2C12 on HAG-2~HAG-3, HAG-3~HAG-4 and HAG-3. Error bars, mean ± s.e.m. Ordinary one-way ANOVA was used for analysis of the data; n=12 cells. c) Immunofluorescence images of YAP in NIH-3T3 on HAG-2, HAG-3 and HAG-4. Scale bar=30 µm. d) Quantitative analysis of YAP nuclear localization in NIH-3T3 on HAG-2, HAG-3 and HAG-4. Error bars, mean ± s.e.m. Ordinary one-way ANOVA was used for analysis of the data; n=15 cells. e) Immunofluorescence images of p-FAK (Green) and F-actin (Red) in NIH-3T3 on HAG-2, HAG-3 and HAG-4. Scale bar=30 µm. f) Quantitative analysis of p-FAK mean fluorescence intensity (MFI) in NIH-3T3 on HAG-2, HAG-3 and HAG-4. Error bars, mean ± s.e.m. Ordinary one-way ANOVA was used for analysis of the data; n=15 cells. g) Enlarged image of immunofluorescence images of p-FAK in e). Scale bar=5 µm. h) Quantitative analysis of the mean area (MA) of FAs in NIH-3T3 on HAG-2, HAG-3 and HAG-4. Error bars, mean ± s.e.m. Ordinary one-way ANOVA was used for analysis of the dat; n=15 cells.

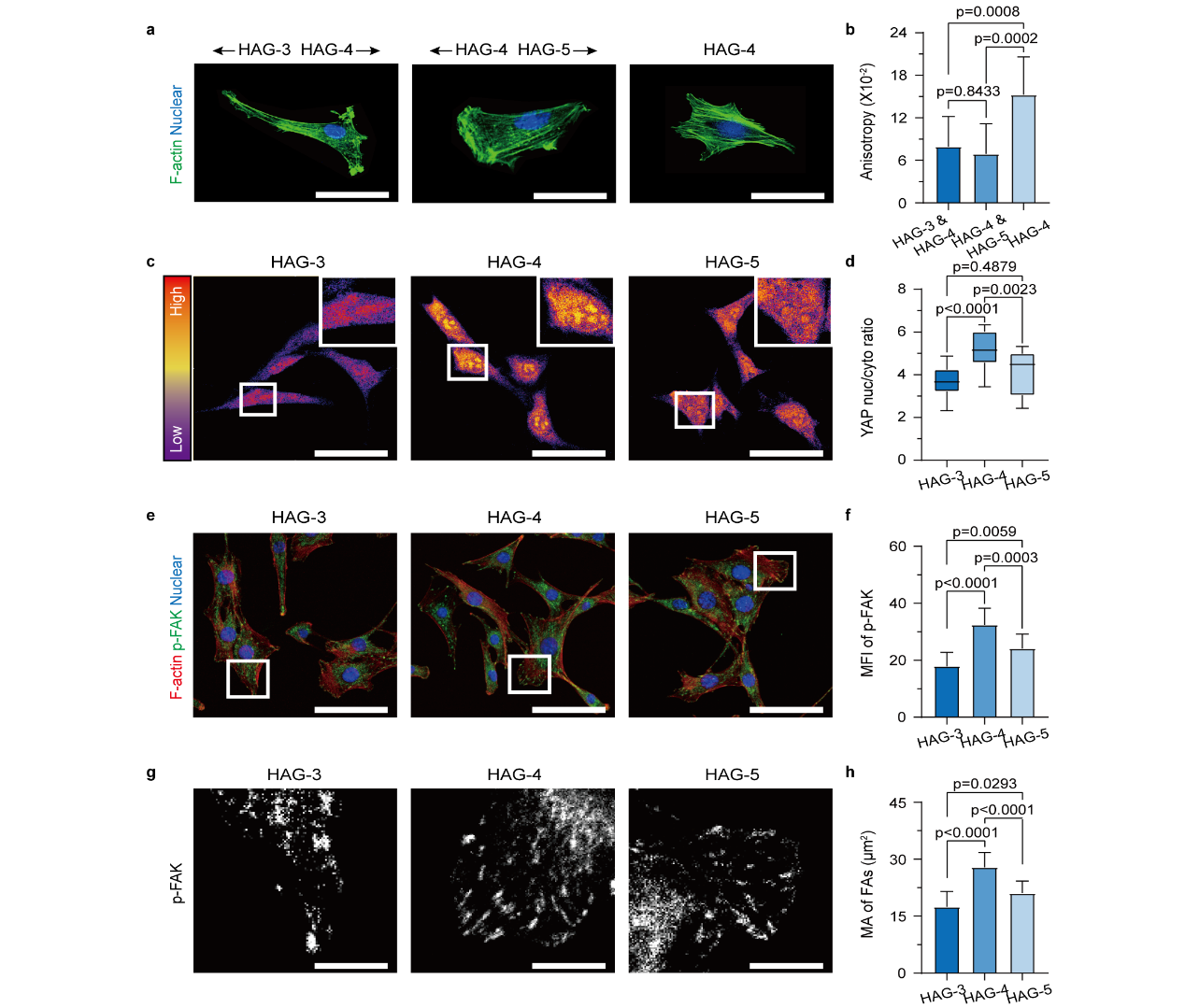

**Supplementary Fig. 14 a)** Immunofluorescence images of F-actin (Green) in chondrocytes on HAG-3~HAG-4, HAG-4~HAG-5 and HAG-4. Scale bar=15 µm. b) Quantitative analysis of anisotropy in chondrocytes on HAG-3~HAG-4, HAG-4~HAG-5 and HAG-4. Error bars, mean ± s.e.m. Ordinary one-way ANOVA was used for analysis of the data; n=12 cells. c) Immunofluorescence images of YAP in chondrocytes on HAG-3, HAG-4 and HAG-5. Scale bar=30 µm. d) Quantitative analysis of YAP nuclear localization in chondrocytes on HAG-3, HAG-4 and HAG-5. Error bars, mean ± s.e.m. Ordinary one-way ANOVA was used for analysis of the data; n=15 cells. e) Immunofluorescence images of p-FAK (Green) and F-actin (Red) in chondrocytes on HAG-3, HAG-4 and HAG-5. Scale bar=30 µm. f) Quantitative analysis of p-FAK mean fluorescence intensity (MFI) in chondrocytes on HAG-3, HAG-4 and HAG-5. Error bars, mean ± s.e.m. Ordinary one-way ANOVA was used for analysis of the data; n=15 cells. g) Enlarged image of immunofluorescence images of p-FAK in e). Scale bar=5 µm. h) Quantitative analysis of the mean area (MA) of FAs in chondrocytes on HAG-3, HAG-4 and HAG-5. Error bars, mean ± s.e.m. Ordinary one-way ANOVA was used for analysis of the data; n=15 cells.

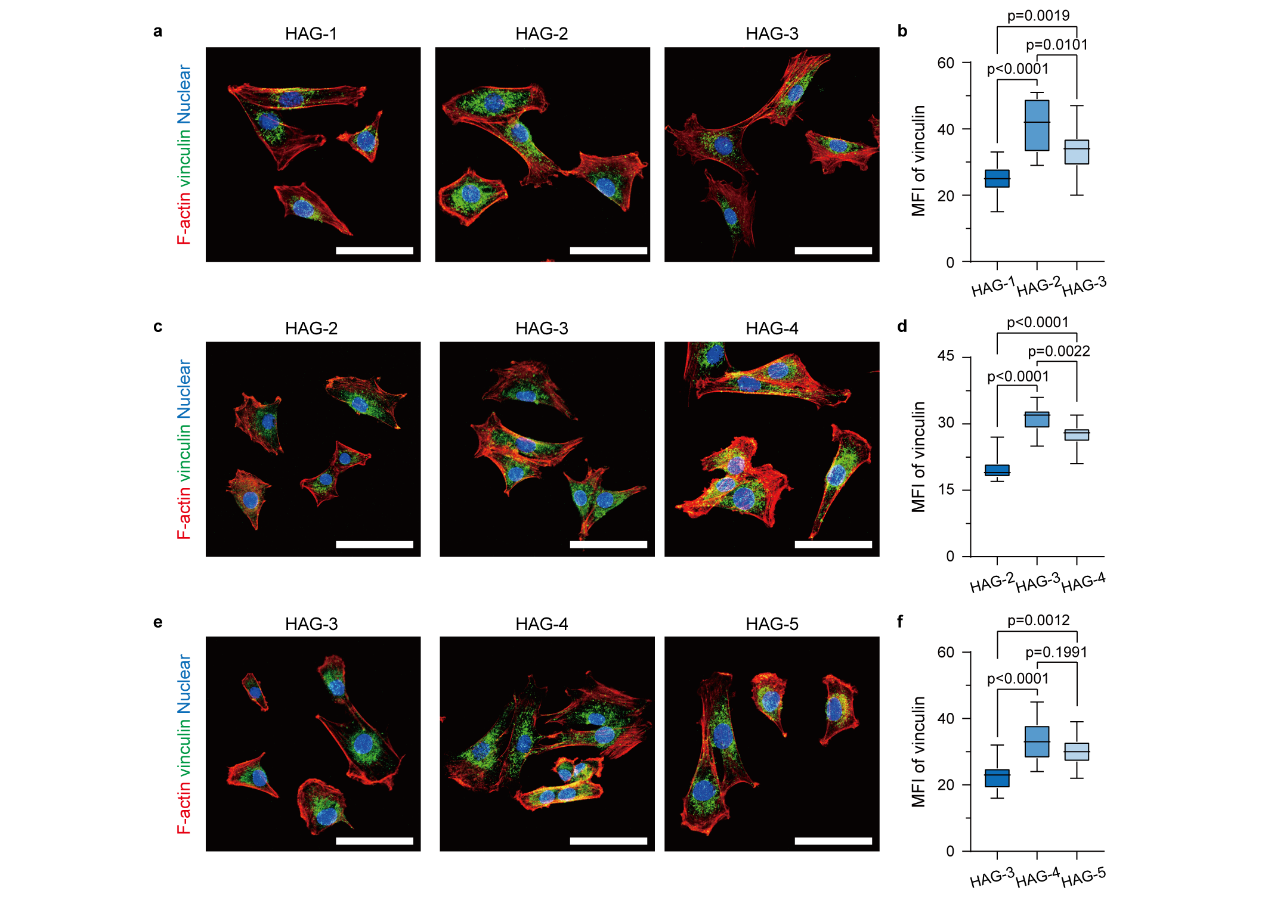

**Supplementary Fig. 15** Immunofluorescence images of vinculin (Green) and F-actin (Red) in a) C2C12 on HAG-1, HAG-2 and HAG-3. Scale bar=40 µm. c) NIH-3T3 on HAG-2, HAG-3 and HAG-4. Scale bar=40 µm. and e) chondrocyte on HAG-3 HAG-4 and HAG-5. Scale bar=40 µm. Quantitative analysis of vinculin mean fluorescence intensity (MFI) in b) C2C12 on HAG-1, HAG-2 and HAG-3. Error bars, mean ± s.e.m. Ordinary one-way ANOVA was used for analysis of the data; n=15 cells. d) NIH-3T3 on HAG-2, HAG-3 and HAG-4. Error bars, mean ± s.e.m. Ordinary one-way ANOVA was used for analysis of the data; n=15 cells and f) chondrocyte on HAG-3 HAG-4 and HAG-5. Error bars, mean ± s.e.m. Ordinary one-way ANOVA was used for analysis of the data; n=15 cells.

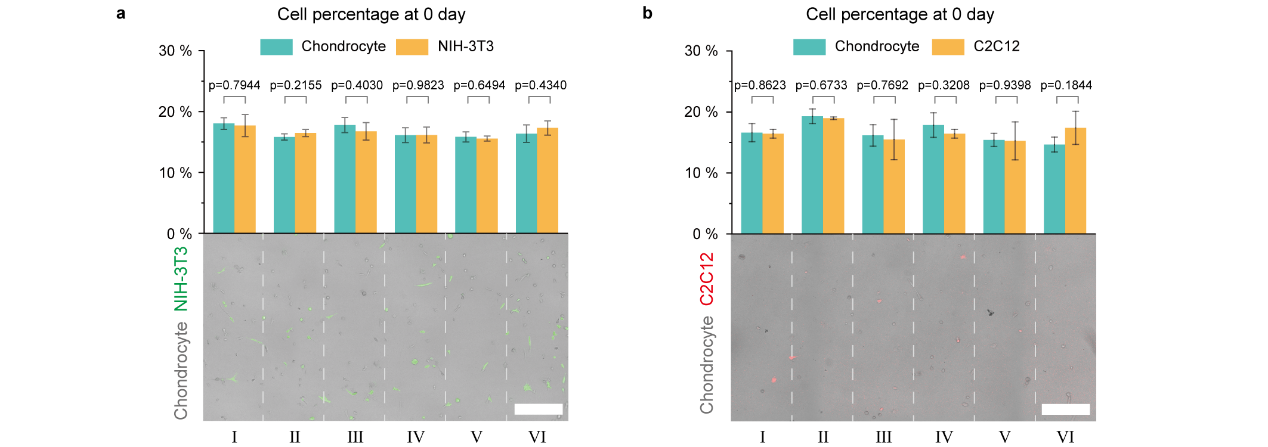

**Supplementary Fig. 16** a) Fluorescent image and analysis of cell percentage (number of A cells in the area/total number of A cells in the figure) of chondrocyte (grey) and NIH-3T3 (green) seeded on HAG-b at day 0. Error bars, mean ± s.e.m. Analysed by the Multiple t tests – one per row. Scale bar = 100 µm. b) Fluorescent image and analysis of cell percentage of chondrocyte (grey) and C2C12 (red) seeded on HAG-c at day 0. Error bars, mean ± s.e.m. Analysed by the Multiple t tests – one per row. Scale bar = 100 µm.
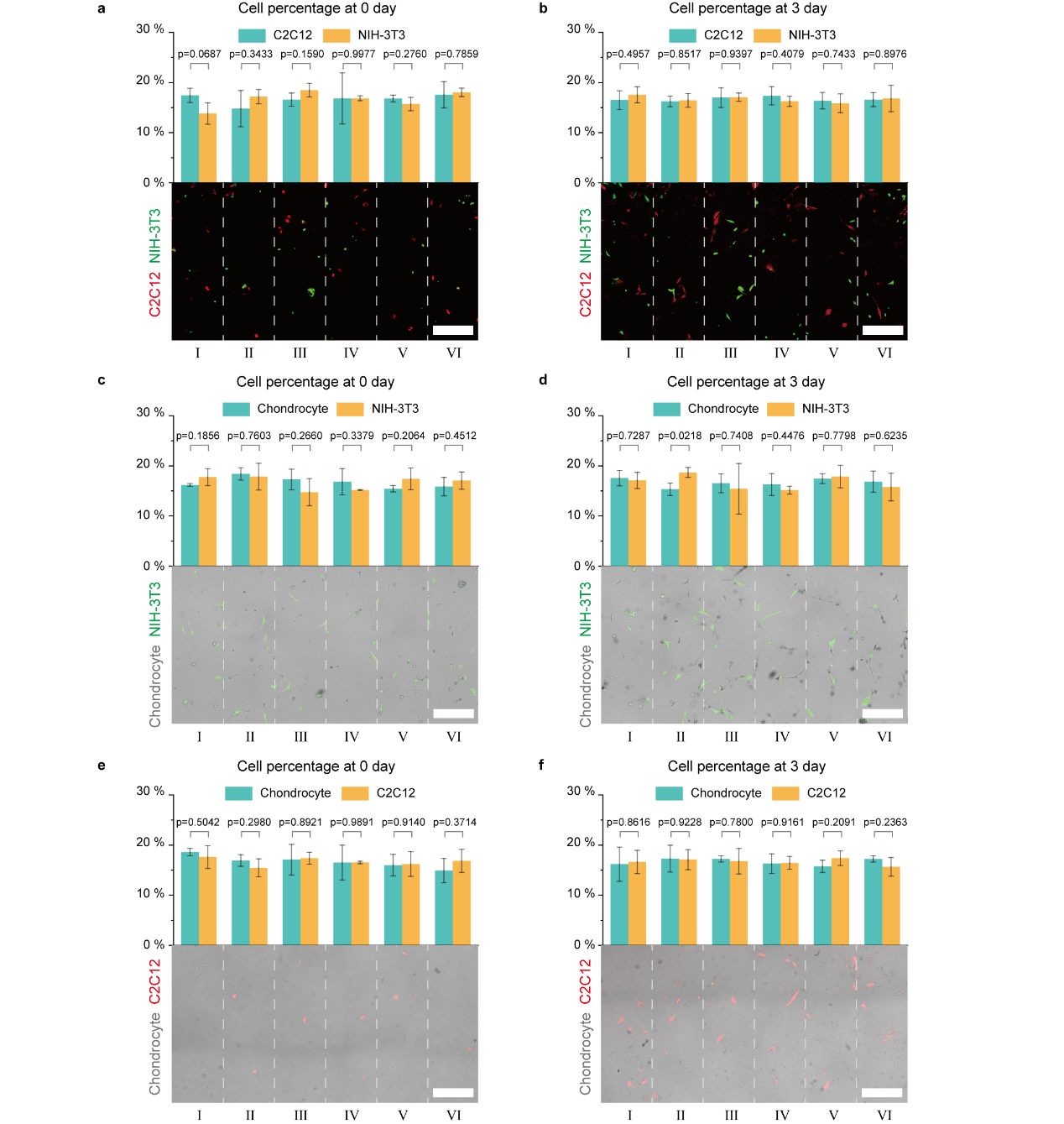

**Supplementary Fig. 17** Fluorescent image and analysis of cell percentage (number of A cells in the area/total number of A cells in the figure) of C2C12 (red) and NIH-3T3 (green) seeded on HAG-2 at a) day 0 and b) day 3. Error bars, mean ± s.e.m. Analysed by the Multiple t tests – one per row. Scale bar = 100 µm. Fluorescent image and analysis of cell percentage of chondrocyte (grey) and NIH-3T3 (green) seeded on HAG-3 at c) day 0 and d) day 3. Error bars, mean ± s.e.m. Analysed by the Multiple t tests – one per row. Scale bar = 100 µm. Fluorescent image and analysis of cell percentage of chondrocyte (grey) and C2C12 (red) seeded on HAG-4 at e) day 0 and f) day 3. Error bars, mean ± s.e.m. Analysed by the Multiple t tests – one per row. Scale bar = 100 µm.

**Supplementary Table 1** Mechanical properties of investigated HAG, were ω is the weight of hydrogel component.

| ω_oxi-HA_ /ω_gelatin_ | Mass fraction (%) | G’  (Pa) | τ_1/2_ ^a^  (s) | ω_oxi-HA_ /ω_gelatin_ | Mass fraction (%) | G’  (Pa) | τ_1/2_ ^a^  (s) |
| --- | --- | --- | --- | --- | --- | --- | --- |
| 0.033 | 10 | 147 | 20 | 0.5 | 10 | 755 | 253 |
|  | 13 | 280 | 126 |  | 13 | 880 | 308 |
|  | 15 | 463 | 184 |  | 15 | 1580 | 386 |
|  | 17 | 710 | 251 |  | 17 | 1450 | 364 |
|  | 20 | 1210 | 356 |  | 20 | 3140 | 456 |
| 0.1 | 10 | 320 | 178 | 1 | 10 | 427 | 182 |
|  | 13 | 477 | 188 |  | 13 | 540 | 197 |
|  | 15 | 813 | 282 |  | 15 | 921 | 310 |
|  | 17 | 1060 | 354 |  | 17 | 1280 | 358 |
|  | 20 | 1540 | 380 |  | 20 | 1750 | 390 |
| 0.2 | 10 | 460 | 191 | 2 | 10 | 175 | 22 |
|  | 13 | 646 | 209 |  | 13 | 220 | 83 |
|  | 15 | 1100 | 362 |  | 15 | 450 | 183 |
|  | 17 | 2200 | 397 |  | 17 | 750 | 253 |
|  | 20 | 2570 | 425 |  | 20 | 1100 | 350 |
